# Loss of TDP-43 drives premature aging and impairs skeletal muscle stem cell pool restoration

**DOI:** 10.64898/2026.08.02.742226

**Authors:** O Pikatza-Menoio, HH Sutcu, M Rodríguez-Hidalgo, A Elicegui, A Vidal-Gil, M Levchuk, N Hernández-Montalvo, L Moreno-Martínez, JM Brito-Armas, R Osta, A Acevedo-Arozena, A López de Munain, S Alonso-Martín

**Author notes:** These authors have equally contributed (HH Sutcu, M Rodríguez-Hidalgo).

## Abstract

TAR DNA-binding protein 43 (TDP-43) dysfunction is a hallmark of amyotrophic lateral sclerosis (ALS) and related disorders, yet its role in skeletal muscle stem cells, the satellite cells (SC), remains incompletely understood. Here, we investigated ALS-associated gain- and loss-of-function TDP-43 mutations together with inducible SC-specific TDP-43 deletion. While *TDP-43^Q331K^* and heterozygous *TDP-43^F210I^* mice displayed normal muscle homeostasis, SC abundance, and regenerative capacity, complete TDP-43 loss caused a marked reduction of the SC pool, particularly in females, and shifted SCs from a CD34^high^ stem-like state toward a CD34^low^ primed population. TDP-43-deficient SCs failed to clonally expand, proliferate, and differentiate, resulting in severe regenerative failure following muscle injury. Notably, the SC pool failed to recover after injury and was nearly depleted 30 days post-injury, accompanied by muscle loss, fibrosis and fat infiltration. Transcriptomic analyses revealed activation of stress and aging-associated programs in uninjured TDP-43-deficient SCs, indicating the premature acquisition of an aging-like state. Consistently, chronological aging further exacerbated SC depletion, establishing TDP-43 as a critical regulator of SC stemness, regeneration, and resistance to age-related decline.

## INTRODUCTION

TAR DNA-binding protein 43 (TDP-43), encoded by the *TARDBP* gene, is a ubiquitously expressed member of the heterogeneous nuclear ribonucleoprotein (hnRNP) family. Through its RNA- and DNA-binding domains, TDP-43 regulates multiple aspects of RNA metabolism, including transcription, alternative splicing, polyadenylation, RNA transport, translation and microRNA biogenesis ^1^. Beyond these canonical functions, TDP-43 has also been implicated in the maintenance of genomic stability, regulation of cell cycle progression, and control of cell survival pathways ^2–4^.

Dysfunction of TDP-43 is a defining feature of several neurodegenerative diseases ^1^. More than 69 *TARDBP* mutations have been associated with amyotrophic lateral sclerosis (ALS) ^5–10^, most of which cluster within the aggregation-prone C-terminal low-complexity domain protein ^11,12^. In both familial (fALS) and sporadic ALS (sALS), pathological TDP-43 mislocalisation from the nucleus to the cytoplasm frequently precedes the formation of insoluble, ubiquitinated and hyperphosphorylated inclusions ^10^. Similar TDP-43 pathology is also observed in frontotemporal dementia (FTD), Alzheimer’s disease (AD), Parkinson’s disease (PD) and limbic-predominant age-related TDP-43 encephalopathy (LATE), highlighting a central role of TDP-43 dysfunction in age-associated degenerative disorders ^1,13,14^.

ALS has traditionally been considered a motor neuron (MN)-driven disease in which skeletal muscle degeneration develops secondary to denervation ^15^. However, increasing evidence suggests that skeletal muscle is not merely a passive target of neurodegeneration ^16,17^. Instead, muscle-intrinsic abnormalities can precede or actively contribute to disease progression, supporting a more complex bidirectional communication between muscle and MNs ^18^. Understanding how TDP-43 dysfunction affects muscle-resident cell populations may therefore provide important insights into both muscle biology and neuromuscular disease pathogenesis.

Skeletal muscle possesses a remarkable capacity for regeneration throughout life. This regenerative potential relies primarily on satellite cells (SC), the resident muscle stem cells located between the myofiber sarcolemma and surrounding basal lamina ^19–23^. Following injury, quiescent SCs become activated, proliferate, and generate myogenic progenitors that subsequently differentiate and fuse to repair damaged fibers or generate new myofibers ^22^. Preservation of the SC pool and its regenerative competence is therefore essential for lifelong muscle maintenance. Conversely, aging is associated with progressive SC dysfunction, impaired self-renewal, loss of stemness, and reduced regenerative capacity, ultimately contributing to muscle decline ^24,25^.

Emerging evidence indicates that TDP-43 plays important roles within the myogenic lineage. We recently demonstrated that primary myoblasts derived from ALS patients display impaired proliferation and differentiation accompanied by reduced levels of both TDP-43 and its downstream target FUS ^17^. Moreover, experimental depletion of either TDP-43 or FUS in healthy myoblasts markedly impaired myotube formation ^17^. Consistent with these findings, Vogler and colleagues reported that TDP-43 assembles into transient cytoplasmic ribonucleoprotein condensates, termed myogranules, during myogenic differentiation, suggesting a physiological role for TDP-43 in muscle formation ^26^. Under pathological conditions, however, these structures may fail to resolve and instead accumulate into aggregates ^26^. Despite growing evidence linking TDP-43 to myogenesis, whether TDP-43 is required for the maintenance, stemness and regenerative function of SCs remains poorly understood. Furthermore, it is unclear whether disease-associated alterations in TDP-43 activity are sufficient to impair SC function or whether more profound loss of TDP-43 activity is required. Addressing this question is particularly relevant in light of the age-dependent decline of both TDP-43 homeostasis and muscle regenerative potential.

In the present study, we investigated the consequences of altered TDP-43 function in skeletal muscle stem cells using ALS-associated gain-(GOF) and loss-of-function (LOF) Knock In (KI) *Tardbp* mutant mice [*TDP-43^Q331K^*and *TDP-43^F210I^*, respectively ^27^] together with a SC-specific inducible TDP-43 knockout (cKO) model. We show that while ALS-associated point mutations do not measurably affect SC function or muscle regeneration in adult mice, complete loss of TDP-43 drives depletion of the stem-like SC compartment, impairs proliferation and differentiation, and causes severe regenerative failure following injury, unable to restore the SC pool. Importantly, transcriptomic and phenotypic analyses revealed that TDP-43-deficient SCs acquire features associated with stem cell aging even under homeostatic conditions, and that aging further exacerbates SC depletion.

Together, our findings identify TDP-43 as a critical regulator of SC stemness and regenerative competence and reveal that loss of TDP-43 promotes the premature acquisition of an aging-like state in skeletal muscle stem cells.

## RESULTS

### Adult mice maintain skeletal muscle and satellite cell homeostasis despite TDP-43 alteration

To determine whether disease-associated alterations in TDP-43 activity are sufficient to impair skeletal muscle and SC function *in vivo*, we first examined KI mouse models: the ALS-associated GOF mutant *TDP-43^Q331K^* ^27–29^ and the LOF mutant *TDP-43^F210I^*, which disrupts the RNA recognition motif 2 (RRM2) required for RNA/DNA binding, leading to the expression of a mutant TDP-43 protein with a strongly diminished RNA binding capacity that leads to TDP-43 LOF and the expression of TDP-43 dependent cryptic exon^27^ (**Fig. S1A**). Because adult homozygous *TDP-43^F210I^*mice are not viable ^27,30^, analyses were performed in adult heterozygous animals, which present no overt phenotypes.

Given the selective vulnerability of fast-fatigable motor units and type II fibers in ALS and related neuromuscular disorders ^31–33^, we evaluated the fast-twitch *tibialis anterior* (TA) and slow-twitch soleus muscles. Histological assessment by hematoxylin and eosin (H&E) staining revealed no detectable abnormalities in muscle architecture or fiber size distribution in either homozygous *Tardbp^Q331K/Q331K^*, hereafter referred as *TDP-43^Q331K^,* or heterozygous *Tardbp^F210I/+^*, hereafter referred as *TDP-43^F210I^*, mice compared with their respective wild-type (WT) littermates (**Fig. S1B**). Likewise, Sirius Red (SR) and Oil Red O (ORO) staining showed no evidence of increased fibrosis or intramuscular fat accumulation in either muscle, although a modest reduction in lipid deposition was observed in the TA of *TDP-43^Q331K^* mice (**Fig. S1C-D**). To determine whether altered TDP-43 function affects muscle fiber composition, we quantified myosin heavy chain (MyHC) isoforms corresponding to type I, IIA, IIX and IIB fibers. Neither *TDP-43^Q331K^* nor *TDP-43^F210I^* mice displayed changes in fiber-type distribution or fiber-type-specific size profiles in either TA or soleus muscles (**Fig. S2**).

To determine whether the absence of a muscle phenotype was accompanied by alterations in the SC compartment, SCs were isolated from adult *TDP-43^Q331K^* and *TDP-43^F210I^* mice. Consistent with the preservation of muscle homeostasis, quantification of PAX7+ SCs revealed no significant differences between mutant and WT mice (**Fig. S3A-B**). SCs were then isolated by fluorescence-activated cell sorting (FACS) based on as CD31negCD45negSca1negIntegrin-α7+CD34+ cell membrane markers ^24^, revealing no differences in SC abundance per gram of muscle compared with WT controls (**Fig. S3C**). Furthermore, the clonogenic capacity of SCs was unaffected by either mutation, as mutant-derived cells formed colonies to a similar extent as WT SCs (**Fig. S3D**). To assess whether these mutations induced broader molecular alterations, RNA sequencing was performed on isolated SCs. Principal component analysis (PCA) of log-normalized counts per million revealed no clear segregation between genotypes, indicating that neither mutation produced major transcriptional changes in the overall SC gene expression profile (**Fig. S3E**).

Together, these findings demonstrate that adult mice carrying either the ALS-associated Q331K mutation or the heterozygous F210I mutation maintain normal skeletal muscle architecture, fiber composition, SC abundance, clonogenic potential, and global transcriptional profiles under homeostatic conditions.

### TDP-43 point mutations do not impair regeneration following muscle injury

To challenge the regenerative capacity of SCs in our mutant models, muscle injury was induced by intramuscular cardiotoxin (CTX) injection into the TA and soleus muscles of *TDP-43^Q331K^*and *TDP-43^F210I^* mice (**Fig. S3A**). Skeletal muscle regeneration was subsequently evaluated 30 days post-injury. As expected, CTX injury increased the number of PAX7+ SCs relative to uninjured muscles ^34^. However, no significant differences in SC abundance were observed between mutant and WT mice in either TA or soleus muscles (**Fig. SA-3B**), indicating that neither mutation impairs SC pool re-establishment following regeneration.

To determine whether regeneration was structurally affected, TA and soleus muscles from *TDP-43^Q331K^*and *TDP-43^F210I^* mice were examined by H&E, SR, and ORO staining 30 days after CTX injury. Histological analyses revealed no differences in muscle architecture or fiber size distribution between mutant and WT groups (**Fig. S4A**). Similarly, collagen deposition and intramuscular fat accumulation remained comparable across genotypes (**Fig. S4B-C**). Assessment of muscle fiber composition by MyHC isoform immunolabeling further demonstrated no changes in the relative proportions or size distribution of individual fiber types in regenerated muscles (**Fig. S4D**).

Together with the preserved SC abundance, clonogenic capacity, and transcriptional profiles observed under homeostatic conditions (**Fig. S3**), these findings indicate that neither the GOF mutation (*TDP-43^Q331K^*) nor the heterozygous LOF mutation (*TDP-43^F210I^*) is sufficient to impair SC function or skeletal muscle regeneration in adult mice (**Fig. S4**). Consequently, we next investigated whether complete ablation of TDP-43 in SCs would reveal functions masked by the partial nature of these disease-associated alterations.

### Loss of TDP-43 compromises satellite cell maintenance, expansion and differentiation

To investigate the consequences of complete TDP-43 loss in SCs, we generated an inducible SC-specific cKO mouse by deleting exon 3 of *Tardbp* in PAX7-expressing cells (*Pax7^CreERT2/+^;Tardbp^fl/fl^*) (**Fig. S5A**). Recombination was induced by tamoxifen (TAM) administration (**Fig. 1A**), and efficient deletion was confirmed two months later by the absence of detectable TDP-43 protein in SCs, but present in the fiber myonuclei, from young adult mice (**Fig. S5B-C**). Despite efficient TDP-43 depletion in SCs, adult cKO mice displayed no overt alterations in skeletal muscle homeostasis (**Fig. S6**). Body weight and grip strength motor performance were comparable to WT littermates in both sexes (**Fig. S6A-B**). Moreover, histological analyses revealed no abnormalities in muscle architecture or fiber size distribution (**Fig. S6C**), and neither SR nor ORO staining detected differences in fibrosis or fat accumulation (**Fig. S6D-E**). Quantification of MyHC isoforms further confirmed the absence of changes in fiber-type composition or fiber-type-specific fiber size (**Fig. S6F**). Together, these findings indicate that TDP-43 depletion in SCs does not compromise the maintenance of pre-existing muscle fibers under homeostatic conditions.

**Figure 1.**
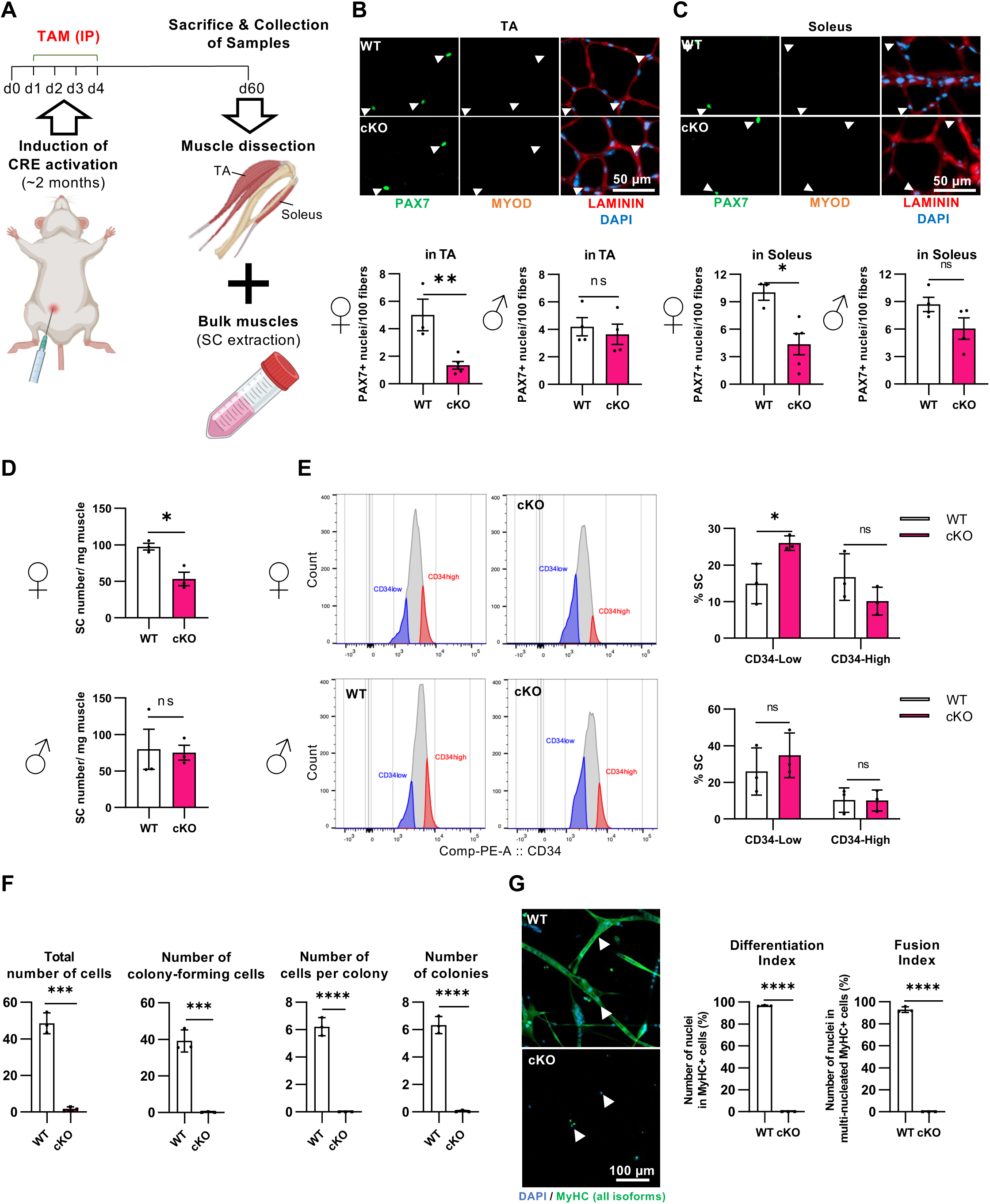
TDP-43 depletion in female satellite cells reduces the pool of stem cells and impairs differentiation. **(A)** Schematics of TDP-43 depletion in satellite cells (SC) by intraperitoneal (IP) tamoxifen (TAM) administration *to Pax7^CreERT2/+^;Tardbp^fl/fl^* (cKO) mice and *Pax7^+/+^;Tardbp^fl/fl^* control (WT) littermates, followed by sample collection. **(B-C)** Upper panels: Representative cross-sectional images of *tibialis anterior* (TA) **(B)** and soleus **(C)** muscles from WT and cKO female mice 60 days (d) post TAM injection, immunolabelled for PAX7 (SCs, green), LAMININ (basal lamina, red) and nuclei with DNA intercalant DAPI (blue). Lower panels: Quantification of PAX7+ SCs in female and male TA **(B)** and soleus **(C)** muscles. Bar graphs represent mean ± SD. N=3-5 mice per condition. **(D)** Number of SCs per mg of processed muscles, immunolabelled for CD34 and ITGA7 markers, isolated by FACS from skeletal muscles of female and male mice. Bar graphs represent mean ± SD. N=3 per condition. **(E)** Quantification of SC subpopulations based on CD34 expression (CD34^low^ and CD34^high^). Bar graphs represent mean ± SD. N=3 per condition. **(F)** Clonogenic potential of FACS-isolated SCs, including total cell number, colony-forming cells, cells per colony and total colonies. Bar graphs represent mean ± SD. N=3 per condition. **(G)** Representative images and quantification of *in vitro* differentiation, assessed by immunolabelled Myosin Heavy chain (MyHC, green) positive myotube formation (arrowheads) and DAPI (blue). Differentiation index indicates nuclei within MyHC+ cells, and fusion index reflects nuclei per myotube. Bar graphs represent mean ± SD. N=3 per condition. Student’s t-test. \**p*<0.05; \*\**p*<0.01; \*\*\**p*<0.001; \*\*\*\**p*<0.0001; ns, not significant.

We next examined whether TDP-43 loss affected the SC pool *in vivo*. Quantification of PAX7+ SCs revealed no significant differences in male mice. In contrast, female mice displayed a pronounced reduction in SC abundance, with a 73% decrease in TA muscles (**Fig. 1B**) and a 54% decrease in soleus muscles (**Fig. 1C**) relative to WT controls. Given this reduction in SC number in female cKO mice, we then asked whether TDP-43 loss may alter SC composition. Previous studies have identified functionally distinct SC subpopulations based on CD34 expression, whereby CD34^high^ (stem-like SCs) cells exhibit greater quiescence, self-renewal, and stemness, whereas CD34^low^ cells are more primed for activation and differentiation ^24^. Consistent with the histological analyses, FACS-sorted SCs from uninjured female cKO muscles exhibited a significant reduction in total SC number compared with WT controls (**Fig. 1D**). Although overall CD34 fluorescence intensity within the SC population was unchanged, subpopulation analysis revealed a 39% reduction in CD34^high^ SCs accompanied by a 1.7-fold increase in CD34^low^ SCs (**Fig. 1E**), indicating depletion of the most stem-like SC compartment and enrichment of the primed population, a feature previously associated with SC aging and loss of regenerative potential ^24^. By contrast, only minor differences were observed in males (**Fig. 1E**). To evaluate the functional consequences of TDP-43 depletion, freshly isolated SCs were subjected to clonogenic assays. WT SCs readily expanded and generated colonies within 72 hours (h), whereas TDP-43-deficient SCs failed to produce detectable colonies (**Fig. 1F**), indicating a profound defect in survival and/or proliferative capacity. Furthermore, during differentiation assays, WT SCs efficiently fused into MyHC+ multinucleated myotubes, whereas cKO-derived SCs failed to undergo myogenic differentiation and showed a marked loss of viable cells over time (**Fig. 1G**).

Together, these findings demonstrate that TDP-43 is required for maintenance of the SC pool and preservation of the stem-like CD34^high^ compartment in female mice. Loss of TDP-43 compromises SC expansion, survival, and differentiation potential, indicating an essential role in sustaining SC function prior to terminal myogenic commitment.

### TDP-43 is essential for satellite cell expansion and differentiation *ex vivo*

Since TDP-43-deficient SCs displayed severe defects in survival, clonogenic expansion, and differentiation *in vitro*, we next sought to assess their behavior within their native niche. To this end, single myofibers were isolated from the *extensor digitorum longus* (EDL) muscles of WT and cKO male or female mice and maintained in culture for up to 72 h (**Fig. 2A**). Under these conditions, quiescent SCs activate, expand, and form clusters along the myofiber surface ^35^. To evaluate SC fate progression, fibers were immunolabeled for PAX7 and MYOD at different time points (**Fig. 2B**). At isolation (0 h), SCs from both WT and cKO mice displayed a quiescent phenotype (PAX7+MYOD-), consistent with the absence of detectable MYOD expression ^36^ (**Fig. 2C**). By 24 h, the majority of SCs had become activated (PAX7+MYOD+) in both genotypes, indicating that TDP-43 depletion does not prevent the initial activation response (**Fig. 2C**). Interestingly, no major sex-dependent differences were observed during the early activation phase. After 48 h in culture, most SCs remained PAX7+MYOD+, although a subset had already lost PAX7 expression while retaining MYOD (PAX7-MYOD+) (**Fig. 2C**). This population was more prominent in cKO fibers, reaching approximately 10% of total cells. Rather than reflecting normal progression through myogenesis, this phenotype suggested premature loss of stem cell identity in the absence of TDP-43. By 72 h, a marked increase in the MYOD-only population was observed specifically in female-derived cKO fibers, where these cells represented up to 60% of the population compared with approximately 30% in WT controls (**Fig. 2C**).

**Figure 2.**
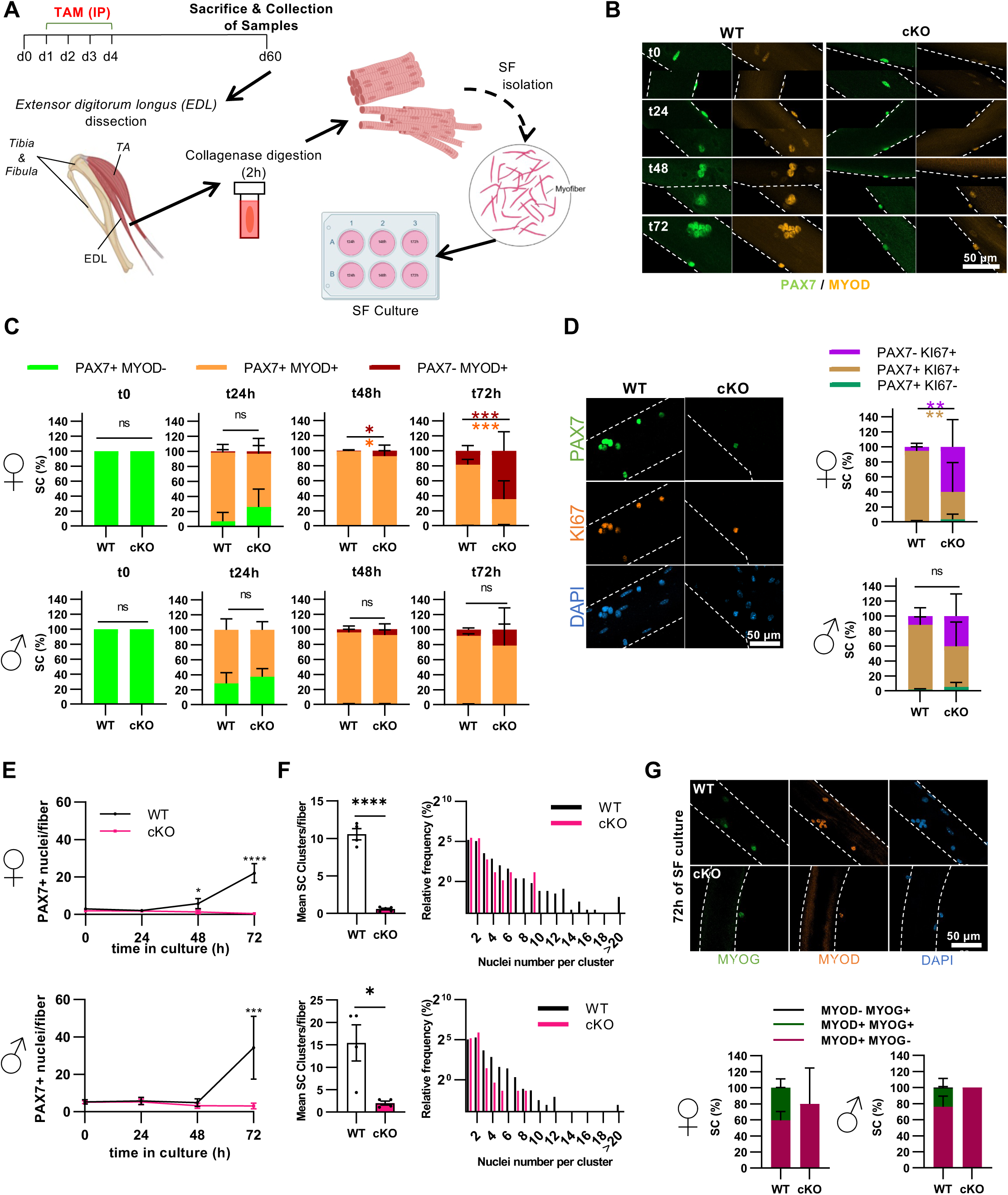
TDP-43 depletion in male and female satellite cells impairs proliferation and differentiation on isolated single myofibers. **(A)** Schematics of TDP-43 depletion in satellite cells (SC) by intraperitoneal (IP) tamoxifen (TAM) administration *to Pax7^CreERT2/+^;Tardbp^fl/fl^* (cKO) mice and *Pax7^+/+^;Tardbp^fl/fl^* control (WT) littermates, followed by *extensor digitorum longus* (EDL) single fiber (SF) isolation and culture. **(B)** Representative images of SCs immunolabelled for PAX7 (green) and MYOD (yellow) on isolated EDL fibers (dotted lines mark myofiber periphery) from cKO male mice and WT littermates over time: 0 (t0) to 72 hours (h) of culture. **(C)** Percentage of SC activation (t24h), proliferation (t48h) and differentiation (t72h) defined by PAX7 and MYOD expression on cultured SF isolated from female (top) and male (bottom) mice. Bar graphs represent mean ± SD. N=4-5 (30-40 fibers per mouse) per condition. **(D)** Representative images (left) and percentage (right) of male SCs immunolabelled with PAX7 and proliferation marker KI67, and nuclei counterstained with DAPI at t72h (dotted lines mark myofiber periphery) from female (top) and male (bottom) mice. Bar graphs represent mean ± SD. N=4-5 (30-40 fibers per mouse) per condition. **(E)** Number of PAX7+ nuclei (SCs) over time per SF isolated from female (top) and male (bottom) mice. Bar graphs represent mean ± SD. N=4-5 (30-40 fibers per mouse) per condition. **(F)** Number (left) and size distribution (right) of SC clusters formed on SFs isolated from female (top) and male (bottom) mice. Bar graphs represent mean ± SD. N=4-5 (30-40 fibers per mouse) per condition. **(G)** Representative images (top) and percentage (bottom) of male SC-derived myogenic cells immunolabelled with MYOD and myogenic differentiation factor MYOGENIN (MYOG), and DAPI at t72h (dotted lines mark myofiber peripheries) from female (left) and male (right) mice. Bar graphs represent mean ± SD. N=2 WT, 5 cKO females; 4 WT, 3 cKO males (30-40 fibers per mouse) per condition. Two-way ANOVA with post-hoc Sidak’s multiple comparison test **(C-E, G)** and Student’s t-test **(F)**. \**p*<0.05; \*\**p*<0.01; \*\*\**p*<0.001; \*\*\*\**p*<0.0001; ns, not significant.

Because MYOD expression alone does not necessarily indicate successful proliferative expansion, we next examined the proliferation marker KI67. Immunolabeling of fibers at 72 h revealed that most remaining cKO-derived cells expressed KI67 (**Fig. 2D**), indicating that TDP-43 deficient SCs retain the capacity to enter the cell cycle. However, these cells also displayed an increased tendency to lose PAX7 expression, consistent with impaired maintenance of stemness. Despite evidence of activation and cell-cycle entry, quantification of SC-derived progeny along the myofiber surface revealed a profound defect in clonal expansion. While WT SCs generated an average of 22 and 34 cells per fiber in female- and male-derived cultures, respectively, repopulation was nearly abolished in cKO fibers from both sexes (**Fig. 2E**). Furthermore, clusters formed by cKO-derived SCs rarely exceeded eight cells, whereas WT SCs generated clusters containing more than 20 cells (**Fig. 2F**). To determine whether these cells could initiate terminal differentiation, fibers were co-immunolabeled for MYOD and Myogenin (MYOG) after 72 h in culture (**Fig. 2G**). Approximately 20% of WT SC-derived cells expressed MYOG, indicating commitment to differentiation. In contrast, MYOG+ cells were virtually absent in cKO-derived clusters, demonstrating a complete failure to initiate the differentiation program (**Fig. 2G**).

Together, these findings demonstrate that TDP-43 is dispensable for the initial activation of SCs but is required for subsequent expansion and differentiation, particularly in female mice. In the absence of TDP-43, SCs prematurely lose stem cell identity, fail to clonally expand, and are unable to initiate the myogenic differentiation program despite entering an activated state.

### Loss of TDP-43 in satellite cells severely compromises muscle regeneration and stem cell restoration

To determine whether complete loss of TDP-43 affects SC-mediated regeneration *in vivo*, muscle injury was induced 30 days after tamoxifen administration by intramuscular cardiotoxin CTX injection into the gastrocnemius, TA, EDL, and soleus muscles (**Fig. 3A**). Muscle regeneration was subsequently assessed at 7- and 30-days post-injury. Consistent with the absence of overt defects under homeostatic conditions, TDP-43 cKO mice displayed normal muscle morphology prior to injury (**Fig. S6**). In striking contrast, 30 days after CTX injury, regeneration was profoundly impaired in cKO mice (**Fig. 3 and S7**). Notably, despite the greater baseline depletion of SCs in female cKO mice, regenerative failure was comparably severe in both sexes following injury. Injured gastrocnemius, TA, EDL, and soleus muscles exhibited a marked reduction in muscle mass, whereas the non-injured quadriceps remained comparable to WT controls (**Fig. 3B and S7A**). Despite cKO mice displaying decreased masses of injured hindlimb muscles, gripping force values recorded prior CTX injection and upon sacrifice at post-injury days 7, 14, 21, and 30 did not reveal differences between cKO and WT mice (**Fig. 3C and S7B**). In line with decreased muscle weights, histological analysis further revealed a severe loss of regenerated muscle tissue, with a dramatic reduction in the number of remaining fibers per area compared with WT muscles (**Fig. 3E**). This regenerative failure was accompanied by extensive fibrosis and adipogenic replacement, as demonstrated by increased SR and ORO staining, respectively (**Fig. 3F-G**). Besides, embryonic myosin heavy chain (eMyHC) staining revealed the presence of few eMyHC+ fibers in injured muscles from cKO mice, while none were detected for WT (**Fig. 3D**).

**Figure 3.**
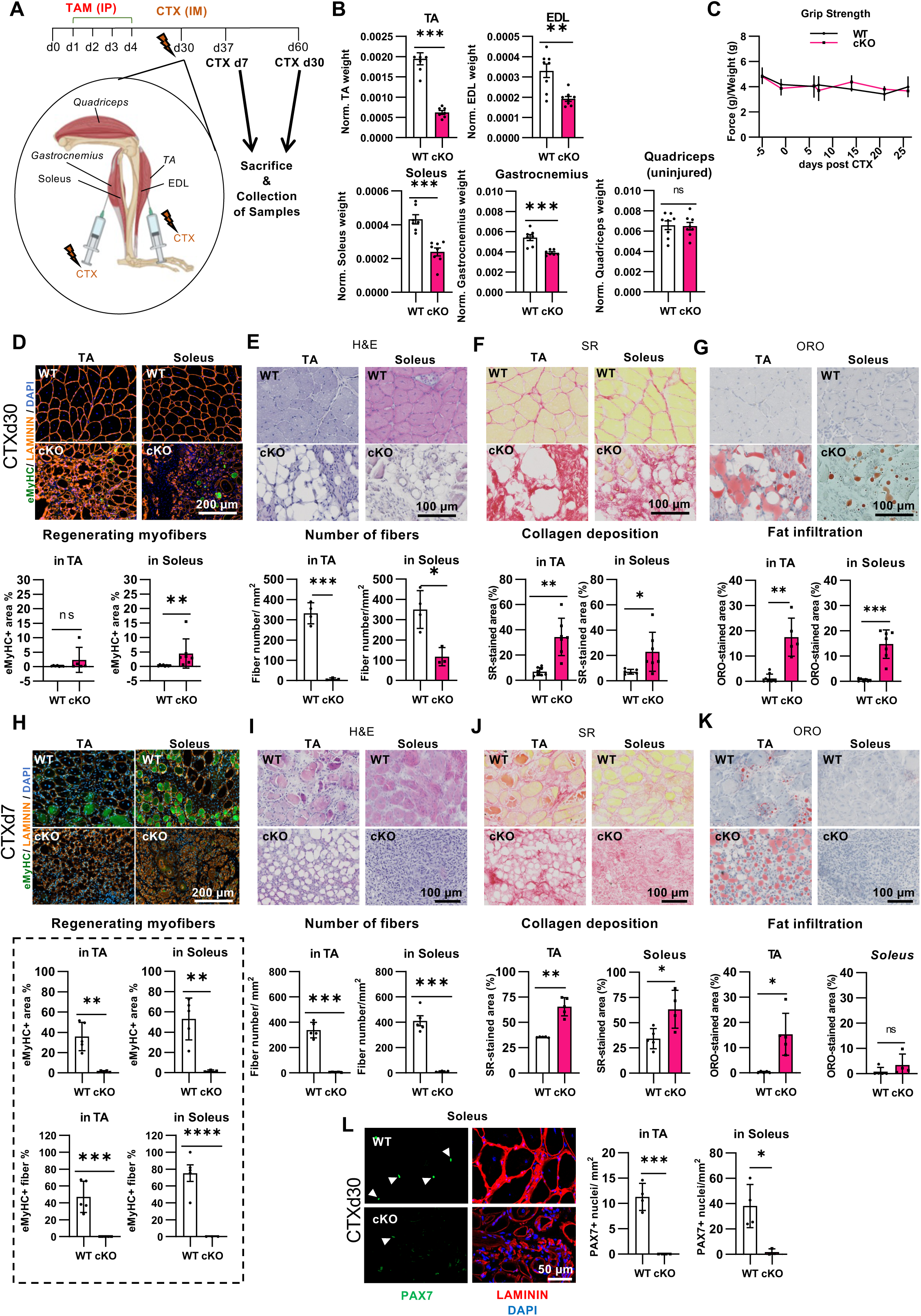
TDP-43 depletion in male satellite cells abolishes muscle regeneration and stem cell pool restoration upon cardiotoxin injury. **(A)** Experimental scheme of TDP-43 depletion in *Pax7*^CreERT2/+^;*Tardbp*^fl/fl^ (cKO) mice and *Pax7*^+/+^;*Tardbp*^fl/fl^ control (WT) littermates by intraperitoneal (IP) tamoxifen (TAM) administration, followed by cardiotoxin (CTX)-induced muscle injury by intramuscular (IM) injection. (B) *Tibialis anterior* (TA), soleus, *extensor digitorum longus* (EDL), gastrocnemius and quadriceps muscle weights 30 days (d) post CTX-injury (CTX d30) dissected from male WT and cKO male mice. Bar graphs represent mean ± SD. N= 7-8 WT, 7-8 cKO per condition. **(C)** Grip strength normalized to body weight over d7-14-21-30 time-points post-CTX. **(D-G)** Top: Representative images; bottom: quantifications of TA and soleus muscles at CTX d30 stained with **(D)** embryonic myosin heavy chain (eMyHC, newly formed myofibers, green), LAMININ (basal lamina, orange), and DAPI (nuclei, blue) and quantification of eMyHC+ area; **(E)** Haematoxylin & Eosin (H&E) for morphology and quantification of number of fibers per area; **(F)** Sirius Red (SR) for collagen deposition (area quantified); and **(G)** Oil Red O (ORO) for fat infiltration (area quantified). Bar graphs represent mean ± SD. N=3-7 per condition. **(H-K)** As for **(D-G)**, top: Representative images; bottom: quantification of TA and soleus muscles at CTX d7 stained by **(H)** eMyHC (green), LAMININ (orange), and DAPI (blue); **(I)** H&E for morphology and quantification of number of fibers per area (top) and per fiber %; **(J)** SR; and **(K)** ORO. Bar graphs represent mean ± SD. N=4-5 per condition. **(L)** Left panel: Representative cross-sectional images of soleus muscles from WT and cKO mice at CTX d30 immunolabelled for PAX7 (satellite cells-SC, green), LAMININ (red) and DAPI (blue). Right panel: quantification of PAX7+ SCs in TA and soleus muscles at CTX d30. Arrowheads indicate SCs. Bar graphs represent mean ± SD. N=4. Student’s t test. \**p*<0.05; \*\**p*<0.01; \*\*\**p*<0.001; \*\*\*\**p*<0.0001; ns, not significant.

Because skeletal muscle regeneration is normally largely complete within 28 days following CTX injury ^21^, we next examined an earlier stage of regeneration. Histological analyses performed 7 days after injury revealed that the regenerative defect was already evident at this stage. Expression of eMyHC, a marker of newly regenerated myofibers, was markedly reduced in both TA and soleus muscles from cKO mice (**Fig. 3H and S7C**). Consistent with impaired myofiber formation, regenerating muscles displayed a reduced number of fibers per area (**Fig. 3I and S7D**), increased collagen deposition (**Fig. 3J and S7E**), and enhanced fat infiltration (**Fig. 3K and S7F**), particularly in the TA muscle.

Given the pronounced defects in SC survival, clonogenic expansion, and differentiation observed *in vitro* and *ex vivo*, we next examined whether the SC compartment could be replenished following *in vivo* regeneration. Whereas WT muscles restored the SC pool after injury, TDP-43 deficient muscles failed to do so. By 30 days post-injury, the number of PAX7+ SCs was dramatically reduced and the residual SC pool was nearly exhausted (**Fig. 3L and S7G**), indicating that TDP-43 is required not only for regeneration of damaged muscle but also for re-establishment of the muscle stem cell compartment following injury.

Together, these findings demonstrate that TDP-43 is indispensable for SC-driven muscle regeneration. Complete loss of TDP-43 results in regenerative collapse characterized by failure of myofiber formation, progressive fibrosis and fat replacement, and an inability to replenish the SC pool, ultimately leading to near-complete exhaustion of the regenerative stem cell compartment.

### Loss of TDP-43 induces stress and premature aging-associated transcriptional programs in satellite cells

To investigate the molecular mechanisms underlying the defects in SC maintenance and regeneration caused by TDP-43 depletion, we performed bulk RNA sequencing on FACS-isolated SCs from adult cKO and WT mice. Differential expression analysis revealed extensive transcriptional reprogramming in TDP-43-deficient SCs in both sexes, with 181 upregulated (UR) and 223 downregulated (DR) genes in males, and 194 UR genes and 181 DR genes in females (pAdj < 0.05, |log2FC| > 0.5) (**Fig. 4A-B**). Functional enrichment analyses identified dysregulation of pathways associated with cell-cycle control, TGFβ signalling, mitochondrial function, stress responses, and cellular homeostasis (**Fig. 4C-D**). While some sex-specific differences were evident, including greater enrichment of G2/M-associated pathways in males and p53-, apoptosis-, and PI3K-mTOR-related programmes in females, both sexes converged on common biological processes linked to mitochondrial dysfunction, altered TGFβ signalling, and impaired cellular homeostasis (**Fig. 4C-D**).

**Figure 4.**
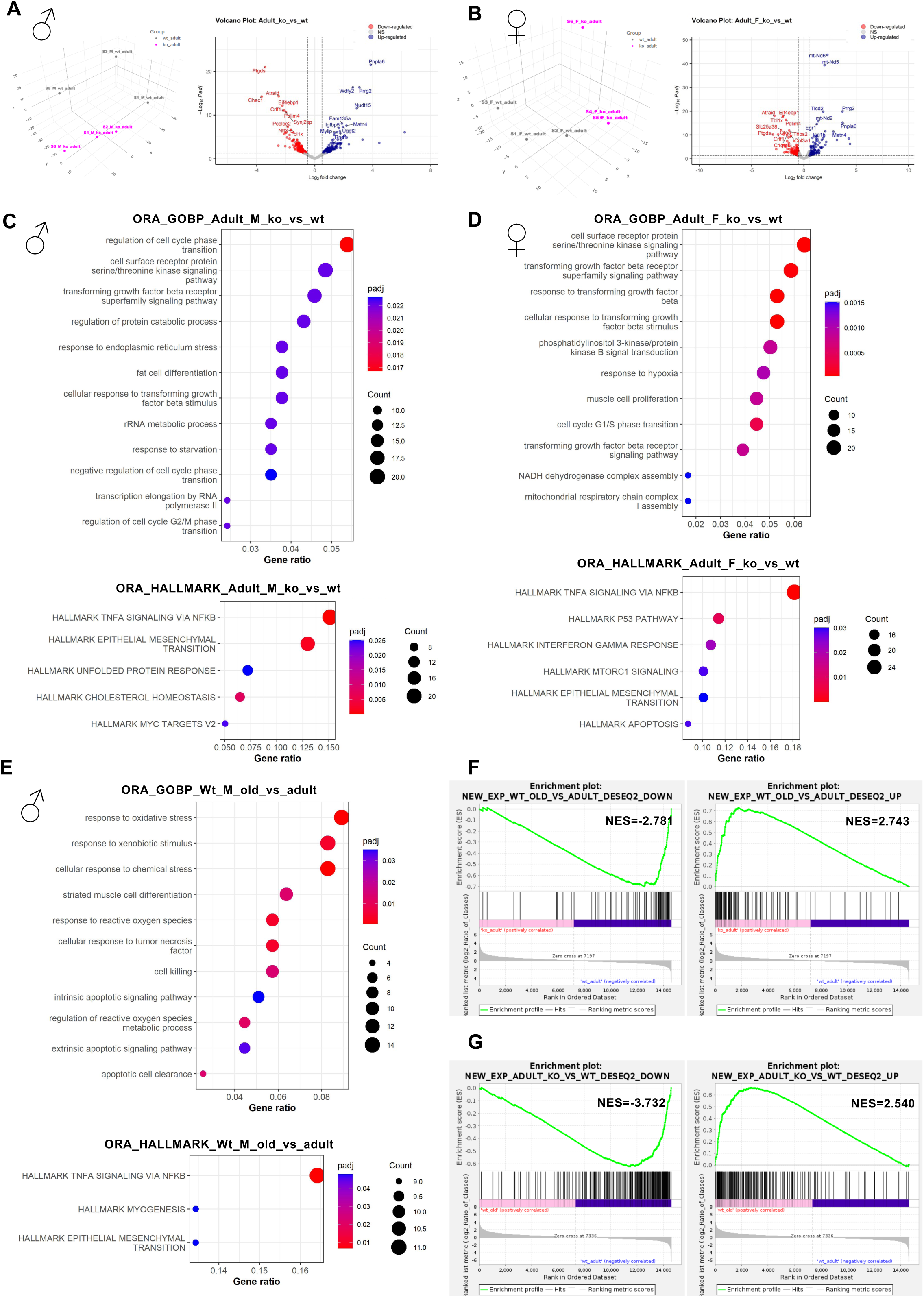
Loss of TDP-43 induces stress and premature aging-associated transcriptional programs in satellite cells. **(A-B)** Principal Component Analysis (PCA) of RNA-seq and Volcano Plot of differentially expressed genes (DEG) (|log₂FC| ≥ 0.5, adjusted *p*<0.05) for satellite cells (SC) from *Pax7*^CreERT2/+^;*Tardbp*^fl/fl^ (cKO) and control *Pax7*^+/+^;*Tardbp*^fl/fl^ control (WT) male **(A)** and female **(B)** mice. **(C-D)** Overrepresentation analyses (ORA) inputting the DEGs against the Gene Ontology Biological Process (GOBP) and Hallmark datasets for SCs from adult male **(C)** and female **(D)** mice. **(E)** ORA inputting the DEGs against the GOBP and Hallmark datasets for SCs from old and adult male mice. **(F)** Gene Set Enrichment Analysis (GSEA) of the WT aging-associated gene signature from RNA-seq data of SCs from old and adult WT male mice over adult cKO and WT male SCs. **(G)** GSEA of the cKO-associated gene signature from RNA-seq data of SCs from adult cKO and WT male mice over old and adult WT male SCs.

To determine whether these transcriptional changes reflected features of physiological aging, we performed RNA-seq analysis of SCs isolated from aged WT mice (**Fig. 4E-G**). As expected, aged SCs displayed enrichment of pathways related to stress adaptation, mitochondrial dysfunction, epithelial-to-mesenchymal transition (EMT), and chronic TGFβ signalling (**Fig. 4E**), all hallmarks of stem cell aging ^37–40^. Notably, several of these pathways were similarly altered in adult cKO SCs, suggesting that loss of TDP-43 induces molecular features normally associated with aged SCs ^41,42^. To directly compare both conditions, Gene Set Enrichment Analysis (GSEA) was performed using aging-associated and cKO-derived transcriptional signatures (**Fig. 4F-G, S8 and S9**). Adult cKO SCs showed significant enrichment for genes associated with physiological aging, whereas aged WT SCs reciprocally displayed enrichment of the cKO signature (**Fig. 4F-G**). In contrast, only minor transcriptomic differences were detected between age cKO and aged WT males (**Fig. 5A-B**), and even fewer changes were observed between adult and aged cKO SCs (**Fig. S8B and S9A**), suggesting that much of the aging-associated transcriptional programme had already been established upon TDP-43 depletion in adulthood.

**Figure 5.**
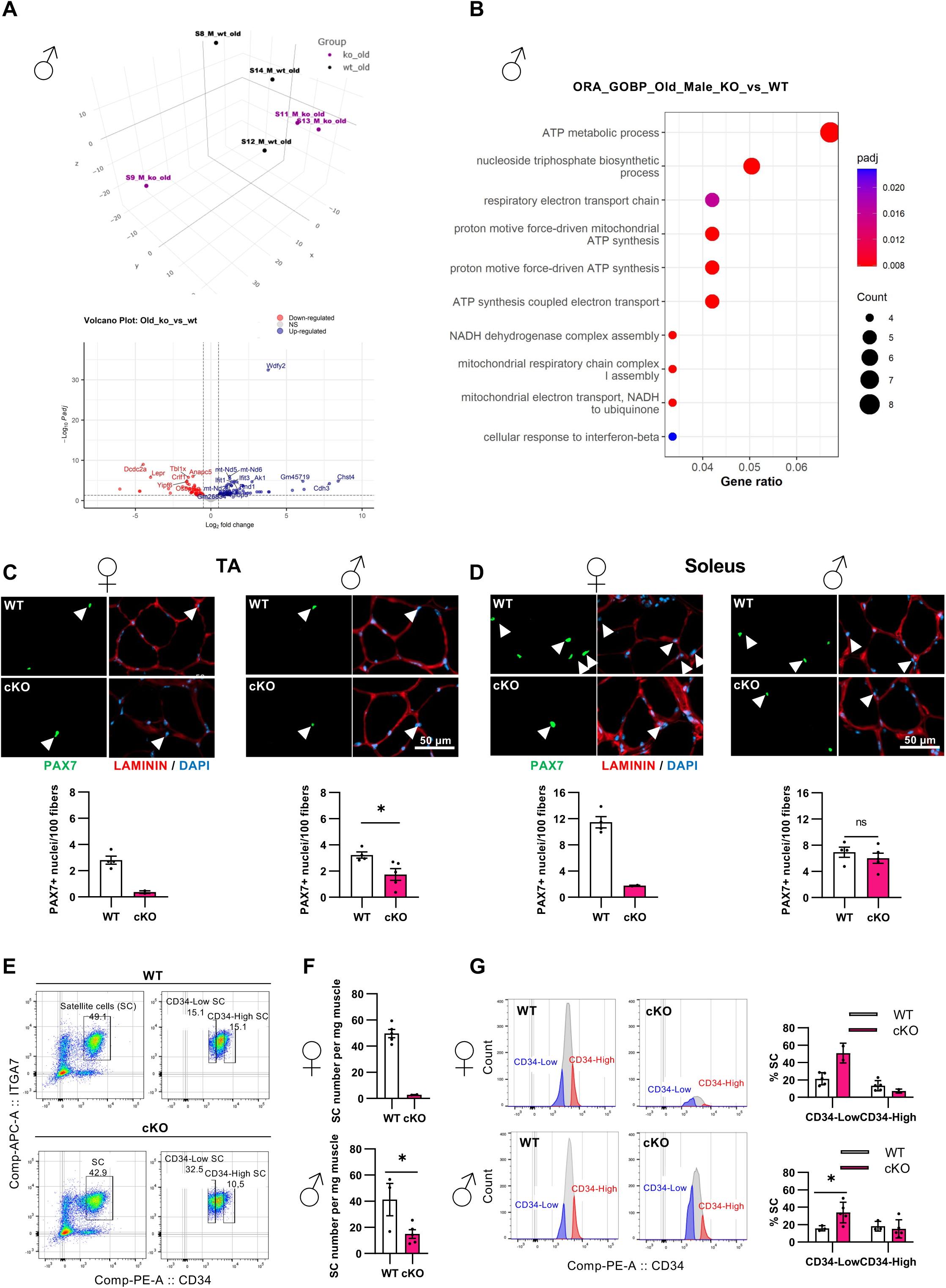
TDP-43 depletion in male and female old satellite cells reduces the pool of stem cells and impairs differentiation. **(A)** Principal Component Analysis (PCA) of RNA-seq and Volcano Plot of differentially expressed genes (DEG) (|log₂FC| ≥ 0.5, adjusted *p*<0.05) for satellite cells (SC) from old *Pax7*^CreERT2/+^;*Tardbp*^fl/fl^ (cKO; N=3) and *Pax7*^+/+^;*Tardbp*^fl/fl^ control (WT; N=4) males. **(B)** Overrepresentation analyses (ORA) inputting the DEGs against the Gene Ontology Biological Process (GOBP). **(C-D)** Representative images and quantification of SCs in cross-sections of *tibialis anterior* (TA) **(C)** and soleus **(D)** muscles immunolabelled with PAX7 (SCs, green), LAMININ (basal lamina, red), and nuclei counter-stained with DAPI (blue) from two-year-old female and male cKO mice and WT littermates. Bar graphs represent mean ± SD. N= 4 WT, 2 cKO female; 4 WT, 5 cKO male mice. **(E-F)** Representative FACS gating strategy used for SC isolation based on CD34 and ITGA7 expression **(E)** and number of SCs isolated per mg of processed muscles **(F)**. Bar graphs represent mean ± SD. N= 5 WT, 2 cKO female; 3 WT, 5 cKO male mice. **(G)** Left: gating strategy; right: quantification of SC subpopulations based on CD34 expression (CD34^low^ and CD34^high^). Bar graphs represent mean ± SD. N= 5 WT, 2 cKO female; 3 WT, 5 cKO male mice. Bar graphs represent mean ± SD. Student’s t test **(C-D, F)** and two-way ANOVA with post-hoc Sidak’s multiple comparison test **(G)**. \**p*<0.05; ns, not significant.

Collectively, these findings demonstrate that loss of TDP-43 induces a sex-modulated but convergent transcriptional program characterized by stress responses, mitochondrial dysfunction, and altered TGFβ signalling. Importantly, the substantial overlap between cKO and physiologically aged WT SC signatures indicates that TDP-43 deficiency drives the premature acquisition of an aging-like state in skeletal muscle stem cells.

### Aging exacerbates satellite cell depletion in TDP-43 cKO mice

Transcriptomic analyses indicated that TDP-43-deficient SCs acquire molecular features associated with physiological aging. Indeed, previous studies have reported age-dependent alterations in TDP-43 expression, including a progressive decrease in skeletal muscle ^43^, a phenomenon that we also previously observed in ALS patient-derived myoblasts, where TDP-43 reduction was further exacerbated under disease conditions ^17^.

To determine whether aging potentiates the effects of TDP-43 depletion, we analysed the SC compartment in two-year-old WT and cKO mice from both sexes. Consistent with the phenotype observed in young adults, aged cKO mice displayed a marked reduction in PAX7+ SCs (**Fig. 5C-D**). However, SC depletion was substantially more pronounced than that observed in adult cKO mice (**Fig. 5C-D vs. Fig. 1B-C**), indicating that aging further compromises maintenance of the TDP-43-deficient stem cell pool in both sexes, although the defect is more pronounced in females. Likewise, FACS analysis following the same gating strategy employed for adult mice revealed a further reduction in SC abundance per gram of muscle in both aged male and female cKO mice (**Fig. 5E-F**). Because TDP-43 depletion in adult female mice reduced as well the stem-like CD34^high^ SC population, we next examined whether aging further altered SC composition. Whereas CD34^high^ SCs remained consistently reduced in cKO mice regardless of age, the CD34^low^ population expanded further in both aged male and female cKO mice, increasing from approximately 1.5-fold in adult mice to 2.5-fold in aged mice relative to WT controls (**Fig. 5G**). These findings indicate that aging amplifies the stem-cell defects caused by TDP-43 loss, further reducing the proportion of stem-like SCs and supporting a premature aging-like phenotype. As mentioned before, comparison of transcriptomic profiles from old and adult cKO SCs revealed minor differences, mostly related to ATP synthesis, indicating that the ‘aging-like’ transcriptomic alterations mediated by TDP-43 loss were not exacerbated with chronological aging (**Fig. 5A-B and S9B**).

Given the progressive deterioration of the SC compartment, we sought to determine whether aging and TDP-43 depletion also affected skeletal muscle maintenance under homeostatic conditions. After verifying no differences in body weight (**Fig. S10A**), neither male nor female cKO mice exhibited any impairment in gripping function over time (**Fig. S10B**), and muscle architecture remained remarkably preserved in both sexes (**Fig. S10C-F**). Indeed, despite the marked reduction in SC abundance, muscle fiber size distribution remained largely unchanged in both WT and cKO mice (**Fig. S10C**). Similarly, no substantial differences in collagen deposition or intramuscular fat accumulation were detected with aging (**Fig. S10D-E**). Finally, aging did not alter muscle fiber-type composition nor fiber-type-specific size distributions in cKO mice compared with age-matched WT controls (**Fig. S10F**).

Together, these findings demonstrate that physiological aging exacerbates the depletion of TDP-43-deficient SCs and further expands the primed CD34^low^ compartment. Combined with the transcriptomic overlap between adult cKO SCs and aged WT SCs, these results support the conclusion that loss of TDP-43 accelerates features of SC aging while mature muscle fibers remain largely preserved under homeostatic conditions.

## DISCUSSION

During this study we have shown that GOF mutation or heterozygote LOF mutation is not enough to impair SCs regenerative capacity or intrinsically affect already differentiated muscle tissue in adult mice. Similar to this, depleting TDP-43 in adult PAX7+ SCs had no impact on the already differentiated myofiber size^26^ or fiber type composition of TA and soleus muscles. However, *Tardbp* exon3 excision in SCs did decrease the SC pool specifically in young adult females, coupled with a reduction in the CD34^high^ population concomitant to an increase in primed CD34^low^ cells. High CD34 expression levels are typically associated with higher stemness and self-renewal potential ^24^. Thus, our findings suggest that TDP-43 loss influences the qualitative composition of the population by shifting cells away from a highly stem-like state, particularly in female mice. Such a change could explain the absence of immediate structural defects in uninjured muscle but may predispose the tissue to impaired regenerative responses upon injury or during aging, when the ability to mobilize a robust stem-like pool is critical. Given the established role of TDP-43 in RNA splicing and stability^1^, its loss may alter transcriptomic programs that sustain the most primitive SC subsets. Accordingly, aging further exacerbated SC depletion and stemness loss in cKO mice relative to their WT littermates, supporting a functional interaction between physiological aging and TDP-43-dependent mechanisms controlling stem cell maintenance.

Further investigations of isolated SCs, either alone or within their microenvironment on isolated single fibers, have shown that TDP-43 is not only critical for myogenic fusion^26^ but is also required during the early phases of myogenic lineage commitment. Consistently, our data indicate that TDP-43 depleted SCs do activate and express MYOD. However, clonogenic and proliferation assays, together with analyses of cluster formation on single myofibers, clearly demonstrate that despite this activation, cKO SCs exhibit severely impaired proliferative capacity. As a result, they are unable to repopulate showing no commitment to differentiation, as evidenced by the absence of the differentiation marker MYOG. In cKO mice, regeneration was profoundly impaired: newly formed myofibers were scarce, while collagen deposition and fat infiltration were markedly increased, reflecting fibrotic remodeling. These findings align with recent reports that TDP-43 is essential for myogenic differentiation and muscle formation ^17,26^, and extend them by demonstrating that its absence in SCs leads to a complete failure of tissue repair *in vivo*. Finally, the stem cell pool is not recovered, indicating the relevance of TDP-43 to preserve it. Together, these findings reveal that TDP-43 is dispensable for maintenance of mature myofibers under homeostatic conditions but is essential for preserving SC stemness and regenerative competence upon muscle damage.

Importantly, many of the dysregulated pathways identified in TDP-43-deficient SCs overlap with hallmark molecular features previously associated with SC aging, including enhanced stress signalling, impaired cellular homeostasis, mitochondrial dysfunction, altered nutrient-sensing pathways, and loss of proteostasis ^25^. Consistent with the known role of TDP-43 in maintaining cellular homeostasis and genomic integrity ^4,44^, disruption of its function may compromise mechanisms required for long-term stem cell maintenance ^45^. In support of this concept, RNA-seq analysis revealed convergent alterations in TGFβ signalling, mitochondrial pathways, and cellular homeostasis, in both male and female cKO SCs. These pathways are recognized hallmarks of aging and contribute to the progressive decline of stem-cell function and regenerative competence^38–41^. Notably, direct comparison with aged WT SCs demonstrated a substantial overlap between the transcriptomic signatures of young adult cKO and physiologically aged SCs. Reciprocal GSEA further confirmed significant enrichment of aging-associated programs in adult cKO SCs, indicating that loss of TDP-43 is sufficient to induce molecular features of aged stem cells despite their young chronological age.

Although male and female SCs exhibited partially distinct transcriptional responses, with males showing greater alterations in proteostasis-and stress-associated pathways and females displaying stronger dysregulation of p53, apoptosis, and metabolic signalling networks, both sexes converged on a common aging-associated transcriptional state. Given that these analyses were performed in quiescent SCs, these enrichments likely reflect alterations in the regulatory networks that preserve stem-cell integrity and activation competence rather than active cell-cycle progression ^46,47^. Importantly, this aging-like transcriptomic profile is strongly supported by the functional phenotype of TDP-43-deficient SCs. Depletion of the CD34^high^ stem-like compartment, reduced clonogenic potential, impaired muscle regeneration, and the failure to restore the SC pool following injury collectively resemble key features of aged muscle stem cells. Moreover, the observation that aged cKO SCs exhibited relatively limited additional transcriptional changes compared with adult cKO SCs suggests that much of the aging-associated molecular programme is already established following TDP-43 depletion. Together, these findings support a model in which TDP-43 safeguards SC stemness and regenerative competence by preventing the premature acquisition of an aging-like state.

These results also bear relevance to human disease. As previously indicated, TDP-43 pathology is a hallmark of ALS, inclusion body myositis and several neurodegenerative disorders ^14,48^. Our findings suggest that, in addition to neuronal dysfunction, impaired TDP-43 activity within muscle stem cells may directly compromise regenerative capacity and contribute to progressive muscle deterioration. These observations highlight SC dysfunction as a previously underappreciated component of the broader TDP-43 disease spectrum.

In summary, our findings demonstrate that TDP-43 safeguards SC stemness and regenerative competence while protecting the muscle stem cell compartment from premature aging. Although dispensable for maintenance of mature myofibers under homeostatic conditions, TDP-43 is essential for SC expansion, differentiation, and restoration of the stem cell pool following injury. Future studies defining how TDP-43 regulates aging-associated pathways in SCs may provide new therapeutic opportunities for muscle degeneration and TDP-43-associated diseases.

## Materials and Methods

### Mouse models

The *TDP-43^Q331K^* KI line was generated by CRISPR–Cas9 mutagenesis through C57BL/6J zygote pronuclear injection of CRISPR reagents, as previously described ^49^, and maintained on C57BL/6J background ^27^. Experimental *TDP-43^Q331K^* mice were provided by Dr. Abraham Acevedo-Arozena (University Hospital from the Canary Islands). The *TDP-43^F210I^* (RRM2 mutant) line was originated through ENU mutagenesis screens ^50,51^ and was rederived by *in vitro* fertilization (IVF) using frozen sperm sourced from RIKEN [B6(D2)-Tardbp<RGSC02268> (RBRC10875)] at the CNB Mouse Embryo Cryopreservation Facility (Centro Nacional de Biotecnología, Madrid) followed by ≥5 generations of backcrossing to C57BL/6J to remove passenger mutations. As homozygous F210I mutated animals die during development, *TDP-43^F210I^* heterozygote mice were used in this work.

*Pax7^CreERT2/+^;Tardbp^flfl^* mice were generated by crossing B6.Cg-*Pax7tm1(cre/ERT2)Gaka*/J (JAX#017763, originally deposited by Gabrielle Kardon) with B6(SJL)-*Tardbptm1.1Pcw*/J (JAX#017591). Cre-mediated recombination deletes exon 3 of *Tardbp* selectively in PAX7+ cells after tamoxifen (TAM) administration. Non-Cre carrier littermates, TAM-injected, were used as controls in all experiments.

All mice were genotyped by PCR/qPCR from ear biopsies using published assays: ^27^, ‘Protocol 20897 - Pax7<tm1(cre/ERT2)Gaka>-alternate1’ (The Jackson Laboratory), and ‘Protocol 26545 - Tardbp<tm1.1Pcw>’ (The Jackson Laboratory). Both sexes were used as indicated in figure legends. For the purposes of this study, mice aged 4.5-12 months served as the adult group, while those between 23-28-month-old represented the aged.

### Animal care and ethics

Mice were maintained on a 12-hours (h) light/dark cycle with food and water *ad libitum*. *TDP-43^Q331K^* (homoeostasis and injury experiments) and *TDP-43^F210I^* (homeostasis), were continuously housed at the animal facility at Universidad de La Laguna (Tenerife, Spain) where all the mice were maintained according to the institutional guidelines of the SEGAI-University of La Laguna and the Government of Canarias, under the regulation CEIBA2016-0194. *TDP-43^F210I^* animals used for injury experiment were initially hosted in Specific Pathogen Free (SPF) facility at Biobizkaia Health and Research Institute (HRI) under the regulation OEBA-CET-2021-004. Following weaning, they were transferred to the SPF facility at Biogipuzkoa HRI. For all models including the *Pax7^CreERT2/+^;Tardbp^flfl^* mouse line generated at or transferred to Biogipuzkoa HRI, procedures were approved by the local institutional committees of Biogipuzkoa HRI and Diputación Foral de Gipuzkoa, under the project licenses PRO-AE-SS-193 (OH-20-37) and PRO-AE-SS-235 (OH21-42).

### Tamoxifen-induced recombination

In *Pax7^CreERT2/+;^Tardbp^fl/fl^* mice, PAX7+ cell-specific TDP43 depletion were achieved by intraperitoneal (IP) injections of 2 mg TAM dissolved in corn oil (20 mg/mL). Animals received 100µL (75-80 mg TAM per kg of body weight) per day for four consecutive days. Analyses were performed 30-60 days after the first injection to ensure efficient recombination.

### Cardiotoxin-induced muscular injury

Before injury, subcutaneous meloxicam (0.2 mg/kg) was administered for analgesia at the time of the procedure. Inhalation anaesthesia was induced using 4% isoflurane and 1.5% oxygen and maintained throughout the procedure with 3% isoflurane and 1.5% oxygen. To induce muscle injury, localized intramuscular injection (IM) of 10 µM cardiotoxin (CTX, *Naja pallida*, Latoxan #L8102) in saline solution was performed using a bevelled 29G needle. Targeted muscles included the fast-twitch *tibialis anterior* (TA), *extensor digitorum longus* (EDL), gastrocnemius, and the slow-twitch soleus muscles. The maximum injection volume was 0.125 mL/kg per point; specifically, 50 µL per TA (simultaneously injure the underlying EDL), and to target both the gastrocnemius and soleus, three-point injections of 25 µL were performed. Muscles were dissected for analyses at 7 days, 14 days, 21 days, and 30 days post CTX injury (referred as CTX d7, CTX d14, CTX d21 and CTX d30, respectively).

### Grip strength test

To assess motor performance and quantify muscular strength in mice, grip strength was measured using a grip strength meter (BIOSEB) equipped with a metal grid connected to a force sensor. Mice were held by the tail and gently lowered onto the grid; once they grasped it, they were pulled horizontally until they released their grip. The peak force (g) exerted immediately prior to release was recorded. To minimize variability due to inconsistent performance, each mouse underwent five trials per session. The mean of the three highest values was calculated and normalized to body weight.

### Isolation of satellite cells by Fluorescence-Activated Cell Sorting

Muscles were manually minced, and enzymatically dissociated , as previously described^24^, for 2 h horizontally shaking (60 rpm) on a rocker at 37 °C with dissociation medium consists of 5 mg/mL of Liberase (0.75 mg/g of tissue; Roche) and 0.2% Dispase (Gibco) in high glucose DMEM (Gibco) supplemented with 1% penicillin/streptomycin (P/S), 0.4 µM CaCl_2_ and 5 µM MgCl_2_. After enzymatic dissociation, where the samples were supplemented with fetal bovine serum (FBS) (1/3 final volume) for enzymatic inactivation, the digests were filtered through 100 µm and 70 µm cell strainers (Corning), respectively. The flow throughs were pre-spinned for 10 minutes (min) at 50 g, 4 °C, and collected supernatants were centrifuged at 430 g for 15 min at 4 °C. Cell pellets were subsequently treated with red blood cell lysis buffer (BD Pharm Lyse) for 10 min on ice in the dark, washed with ice-cold PBS and filtered through 40 µm strainers (Corning), followed by another centrifugation step. The final cell pellets were stored in FBS with 10% DMSO at -80 °C and later in liquid nitrogen until used.

To isolate SCs by FACS, thawed pellets were first washed in cold PBS containing 2% goat serum and 1% P/S (FACS buffer). Upon centrifugation at 430 g for 15 min at 4 °C, cell pellets were resuspended in 700 µL of FACS antibody mix prepared in FACS buffer and were incubated for 1 h on ice in the dark. Cells were then washed in FACS buffer, centrifuged as above; pellets were re-suspended in 200 µL FACS buffer. Before sorting, 0.2 µL of DAPI (5 mg/mL) was added for dead-cell exclusion, and the suspension was filtered through a 30 µm Filcon® syringe filter (BD). Sorting was performed using a FACS Aria IIu (BD) equipped with a 70 µm nozzle and set to 4-way purity mode at the flow cytometry platform at Centro de Investigación Médica Aplicada (CIMA, University of Navarre). After gating for DAPI-negative (viable) cells, the SC population was identified as CD34+ITGA7+ using PE-conjugated anti-CD34 and APC-conjugated anti-α7 integrin (ITGA7). Lineage-positive cells were excluded using a PE-Cy7-conjugated “dump” channel containing antibodies against CD45, Sca1, and CD31 (CD45-Sca1-CD31-). Isolated quiescent SCs were either processed for RNA isolation using the RNeasy Qiagen Micro Kit or plated into 96 well (p96) plates for *in vitro* assays. Data acquisition used BD FACSDiva v8.0.2, and downstream analyses were carried out with FlowJo v10.4.

### In vitro satellite cell culture

For all *in vitro* experiments, FACS-sorted SCs were incubated in collagen type I-coated (0.05mg/mL) dishes at 37 °C with 5% CO_2_. Unless indicated, SCs were cultured in Clonogenic Medium (DMEM containing 1% P/S, 4 mM L-glutamine, 10% FBS, 5% horse serum [HS], 100 mM HEPES, 1 mM sodium pyruvate, 2 mM GlutaMax, and 2.5 ng/mL bFGF [Preprotech]). For clonogenic assays, SCs were seeded in low-density (50 cells/well in p96 plates) into regular 96-well plates (Corning), incubated for 72h, and then formed colonies (≥3 cells) were quantified. 8 replicate wells were analysed per biological replicate. For TDP-43 KO verification, SCs were seeded in medium-density (2000 cells/well) and fixed after 72 h in culture. For differentiation assays, cells were seeded to confluency in ibiTreat µ-96 well plates (IBIDI), and 48 h later culture medium was switched to low-serum Differentiation Medium (DMEM containing 1% P/S, 4 mM glutamine and 5% HS) and further incubated for 8 additional days. *In vitro* differentiation capacity of myogenic cells was analysed by MyHC (all isoforms) immunostaining.

### Ex vivo single-fiber culture

Dissected EDL muscles were digested with 0.2% collagenase type I ( Sigma-Aldrich) for 2 h at 37 °C, and intact single myofibers (SF) were subsequently isolated using a fire-polished Pasteur pipet pre-flushed with 0.5% HS-PBS as previously described^52^, and cultured in DMEM containing 1% P/S, 4 mM L-glutamine, 10% HS and 1% chicken embryo extract (MP Bio) in 6-well plates (Corning). SFs were fixed with 4% PFA at 0, 24, 48, and 72 h and stored in 0.1% NaN_3_-PBS at 4 °C. Analysis of SC activation, proliferation, and differentiation on SFs was performed by immunostaining for PAX7, MYOD, KI67, and MYOG and quantification.

### Histology and tissue immunostaining

TA and soleus muscles were embedded in Optimal Cutting Temperature (OCT, Tissue-Tek®) compound, liquid nitrogen cooled-isopentane frozen, cryosectioned (8 µm for samples in homeostasis and injured samples from the *TDP-43^Q331K^* and *TDP-43^F210I^* lines, and 12 µm for injured samples from *Pax7^CreERT2/+^;Tardbp^flfl^* and *Pax7^+/+^;Tardbp^flfl^* mice), and processed for H&E, SR, or ORO, as previously described^17,23^ or immunofluorescence. Fiber-type composition was assessed using MyHC isoform-specific antibodies on unfixed muscle sections. For all the other targets, cryosections were thawed and fixed with 4% PFA, to subsequently permeabilize in 0.5% Triton X-PBS, block with 4% IgG-free BSA (Jackson Immunoresearch), 1% goat serum, 0.025% Tween20-PBS. Primary antibodies were diluted in blocking buffer and incubated in a wet chamber overnight (O/N) at 4 °C. Secondary antibodies and nuclear stain DAPI were diluted in blocking buffer and incubated for 1 h at room temperature (RT). Exceptionally, for TDP-43 immunostaining a fluorophore-conjugated laminin antibody was used as a last step. Samples were washed in 0.025% Tween-20 in PBS and mounted in Fluoromount-G.

### Immunostaining for satellite cell assays performed in vitro and ex vivo

In vitro SC immunostaining was performed by simultaneous permeabilization and blocking in 5% normal goat serum (NGS), 0.3% Triton-X-PBS for 1 h at RT. Primary antibodies were incubated O/N at 4 °C, and after washes, secondary antibodies and DAPI were incubated for 1 h at RT in the dark. SFs were permeabilized in 0.5% Triton X-100 in PBS for 8 min. Following washes in 0.025% Tween-20 in PBS, fibers were blocked for 2 h at RT in blocking buffer containing 10% NGS, 10% normal swine serum (SS) and 0.025% Tween-20 in PBS, with gentle mixing every 20 min. Primary antibodies diluted in blocking buffer were added and incubated overnight at 4 °C on a low-speed rocker. Secondary antibodies and DAPI were diluted in blocking buffer and incubated for 1 h at RT. Fibers were then washed in 0.025% Tween-20 in PBS and mounted onto glass slides in Fluoromount-G. All antibodies used are listed in **Supplementary Tables 1.1 and 1.2**.

### Image acquisition and analysis

For *in vitro* cultured SCs for 72 h, TDP-43 expression was assessed from 10 representative fields per well at 40X magnification using the Axio Observer 7 epifluorescence microscope (Zeiss). For differentiation assays, differentiation and fusion indices were quantified from five representative fields per well at 10X magnification using the LSM 900 confocal microscope (Zeiss). Three replicate wells were analysed per biological replicate.

For tissue analyses, whole-section tiled images were acquired using the ZEISS Axioscan 7 widefield slide scanner, Axio Observer 7 widefield or LSM900 laser scanning confocal microscopes (Carl Zeiss, Inc., Germany). Quantification of SC number, marker expression, fibrosis, fat infiltration, and fiber size was performed blinded, using ImageJ/Fiji v2.16.0. SFs were analysed using the eyepiece in the Axio Observer 7 microscope, and representative images were acquired with the LSM900 laser scanning confocal microscope.

### RNA preparation and RNA-seq performance

Total RNA was extracted from FACS-isolated SCs from at least three independent biological replicates using the RNeasy Micro Kit (QIAGEN), including DNase I treatment. RNA integrity was assessed using the Eukaryote Total RNA Pico kit on the Agilent 2100 Bioanalyzer. Library preparation was carried out using the “NEBNext Single Cell/LowInput RNA library prep Kit for Illumina” from New England Biolabs. NGS experiments were performed in the Genomics Unit of the CNIC; libraries were sequenced on Illumina NextSeq 1000/2000 platform at a sequencing depth of at least 20 M single-end 200 nucleotide long reads (SE200).

### RNA-seq data processing and analysis

The analyses were performed on the Genomic Platform of the Biogipuzkoa HRI using the standardized nf-core/rnaseq (v3.15.0) and nf-core/differential abundance (v1.5.0) pipelines to ensure reproducibility and best practices in computational workflows ^53^. FastQ files were processed through nf-core/rnaseq. The fastp v.0.23.4 tool ^54^ was used for adapter removal and low-quality base filtering (Phred score < 15) applying a sliding window approach. The clean RNA-seq reads were aligned to the *Mus musculus* GRCm38.99 reference genome available on ENSEMBL using the STAR aligner v.2.7.10a ^55^, and the abundance of each transcript was quantified using the Salmon v.1.10.1 tool ^56^. Count matrices generated by nf-core/rnaseq were analysed for differential expression using nf-core/differential abundance. Low-abundance genes were filtered out by retaining genes with a minimum abundance of 10 counts in at least three samples. The DESeq2 R package was used for differential analysis ^57^, adjusting *p*-values using the Benjamini-Hochberg (BH) method. Transcripts were considered differentially expressed if adjusted *p*-value ≤ 0.05 and |log2FC| ≥ 0.5.

Over-representation analysis (ORA) was performed using the R package clusterProfiler v.4.12.6^58^. Specifically, Gene Ontology Biological Processes (GOBP) were analysed using the enrichGO function, Reactome pathways were assessed with the enrichPathway function, and Hallmark gene sets (retrieved via the msigdbr package) were evaluated using the enricher function. Only significant enrichment results (adjusted *p*-value ≤ 0.05) were considered. To evaluate specific transcriptional signatures associated with aging, a baseline aging signature specific to our experimental conditions was established by comparing Wild-Type (WT) Old versus WT Adult samples from our current dataset. Gene Set Enrichment Analysis (GSEA v4.3.2) ^59,60^ was performed using MSigDB collections (v2026.1) ^61,62^ and the aging signatures mentioned above. The analysis was run using a weighted scoring scheme with the log2 ratio of classes as the ranking metric. Significance was assessed using 1,000 gene-set permutations, and gene sets were restricted to those containing between 15 and 500 genes. Significant enrichment was defined by a False Discovery Rate (FDR) q-value ≤ 0.25.

### Statistical analysis

Unless otherwise stated, data were analysed using GraphPad Prism 8. Normality was assessed by Shapiro–Wilk test. Two-group comparisons were performed using an unpaired Student’s *t*-test. For comparisons with more than 2 groups, two-way ANOVA followed by Sidak’s post-hoc multiple comparisons test was used. Results are shown as mean ± SD. Statistical details and sample sizes are provided in figure legends. Where indicated, significance is indicated as \**p*<0.05; \*\**p*<0.01; \*\*\**p*<0.001; \*\*\*\**p*<0.0001.

## Author Contributions

This project was administered by SA-M. SA-M conceived, planned and supervised these experiments. OP-M, HHS, AE, AV-G, ML, NH-M, MM-M, IA conducted laboratory experiments. All data analyses from wet lab experiments were performed by OP-M, and HHS and SA-M reviewed them. JMB-A and AA-A provided mice for the experiments. MR-H performed RNA-seq analysis and interpretation. ALM and SA-M contributed to funding acquisition and supported the project. OP-M, HHS and SA-M wrote, reviewed and edited the original draft. All authors have read, reviewed and approved the final version of this paper.

## Acknowledgments

The authors acknowledge the technical and human support of the Genomics and Animal Facility Platforms of Biogipuzkoa Health Research Institute (HRI; San Sebastian); the Animal Facility of Biobizkaia HRI (Barakaldo) and the Animal facility of Universidad de La Laguna (Tenerife) for hosting the experimental mice used in this study. The authors also thank Dr. Diego O. Alignani and Aitziber López López from the Cytometry Platform at Centre for Applied Medical Research (CIMA, University of Navarra, Pamplona) for their valuable support during the cell sorting experiments. We want finally to thank Dr. Alberto Benguría Filippini from the Genomics Unit at Centro Nacional de Investigaciones Cardiovasculares (CNIC, Madrid) for expertise and assistance in NGS. Finally, authors thank Mikel García Puga for his assistance to deposit the NGS data to GEO.

## Data Availability Statement

RNA-seq data reported in this publication have been deposited in NCBI’s Gene Expression Omnibus (GEO) and are accessible through GEO series accession number GSE339358.

