## Supplementary Tables for "Loss of TDP-43 drives premature aging and impairs skeletal muscle stem cell pool restoration"

**Supplementary Table 1.1. Primary antibodies used in immunofluorescence and Fluorescence-Activated Cell Sorting.**

| Primary Antibodies | Application | Dilution | Host | Company | Reference |
|---|---|---|---|---|---|
| eMyHC | IF | 1:50 | Ms | DSHB | F1.652-s |
| FUS | IF | 1:200 | Rb | Invitrogen | PA5-52610 |
| Ki67 | IF | 1:500 | Rb | Abcam | ab15580 |
| Laminin | IF | 1:300 | Rb | Sigma-Merck | L9393 |
| Laminin AF647 | IF | 1:100 | Rb | Novus Biologicals | NB300-144AF647 |
| MYH1 (MHC all isoforms) | IF | 1:50 | Ms | DSHB | A4.1025-c |
| MYHC I | IF | 2:3 | Ms | DSHB | BA-D5 |
| MyHC IIa | IF | 1:3 | Ms | DSHB | SC-71 |
| MyHC IIb | IF | 1:1 | Ms | DSHB | BF-F3 |
| MyoD1 | IF | 1:100 | Ms | Santa Cruz Biotechnology | sc-377460 |
| Pax7 | IF | 1:50 | Ms | Santa Cruz Biotechnology | sc-81648 |
| TDP-43 | IF | 1:300 | Ms | R&D Systems | MAB7778 |
| AF647 Alpha-7-integrin (R2F2) | FACS | 1 µL / 100 µL ($10^6$ cells) (4 µg/ml final) | Rat | Ablab | 67-0010-05 |
| PE Rat anti-Mouse CD34 (RAM34, RUO) | FACS | 3 µL / 100 µL ($10^6$ cells) (6 µg/ml final) | Rat | BD Biosciences | 551387 |
| PE/Cy7 Rat anti-mouse CD31 (390) | FACS | 0.5 µL / 100 µL ($10^6$ cells) (1 µg/ml final) | Rat | Biolegend | 102418 |
| PE/Cy7 Rat anti-mouse CD45 (30-F11) | FACS | 0.5 µL / 100 µL ($10^6$ cells) (1 µg/ml final) | Rat | Biolegend | 103114 |
| PE-Cy™7 Rat Anti-Mouse Ly-6A/E ScaI (D7, RUO) | FACS | 0.5 µL / 100 µL ($10^6$ cells) (1 µg/ml final) | Rat | BD Biosciences | 558162 |

**Supplementary Table 1.2. Secondary antibodies used in immunofluorescence.**

| Secondary Antibodies | Application | Dilution | Host | Company | Reference |
|---|---|---|---|---|---|
| F(ab')2-Goat anti-Mouse IgG (H+L) Cross-Adsorbed Secondary Antibody, Alexa Fluor™ 488 | IF | 1:500 | Gt | Invitrogen | A-11017 |
| Goat anti-Mouse IgG1 Cross-Adsorbed Secondary Antibody, Alexa Fluor™ 488 | IF | 1:500 | Gt | Invitrogen | A-21121 |
| IgG1 Cross-Adsorbed Goat anti-Mouse, Alexa Fluor™ 647 | IF | 1:500 | Gt | Invitrogen | A-21240 |
| Cy3 AffiniPure Fab Fragment Goat Anti-Mouse IgG2a, Fcγ fragment specific | IF | 1:500 | Gt | Jackson ImmunoResearch Labs | 115-167-186 |
| Alexa Fluor™ 647 AffiniPure Fab Fragment Goat Anti-Mouse IgG2a, Fcγ fragment specific | IF | 1:500 | Gt | Jackson ImmunoResearch Labs | 115-607-186 |
| Cy™3 AffiniPure Goat Anti-Mouse IgG, Fcγ subclass 2b specific (min X Hu, Bov, Rb Sr Prot) | IF | 1:500 | Gt | Jackson ImmunoResearch Labs | 115-165-207 |
| IgM (Heavy chain) Cross-Adsorbed Goat anti-Mouse, Alexa Fluor™ 488 | IF | 1:500 | Gt | Invitrogen | A-21042 |
| F(ab')2-Goat anti-Rabbit IgG (H+L) Cross-Adsorbed Secondary Antibody, Alexa Fluor™ 555 | IF | 1:500 | Gt | Invitrogen | A-21430 |
| F(ab’)2-Goat anti-Rabbit IgG (H+L) Cross-Adsorbed, Alexa Fluor™ 647 | IF | 1:500 | Gt | Invitrogen | A-21246 |
| DAPI (4',6-Diamidino-2-Phenylindole, Dihydrochloride) | IF | 1:1,000 | NA | Invitrogen | 10184322 |
| *IF, immunofluorescence; FACS, Fluorescence-activated cell sorting; Ms, mouse; Rb, Rabbit; Gt, goat; MyHC or MHC, Myosin Heavy Chain; NA: not applicable, (not an antibody) | | | | | |
