## Supplementary Figures and Figure Legends for "Loss of TDP-43 drives premature aging and impairs skeletal muscle stem cell pool restoration"

### Supplementary Figure 1

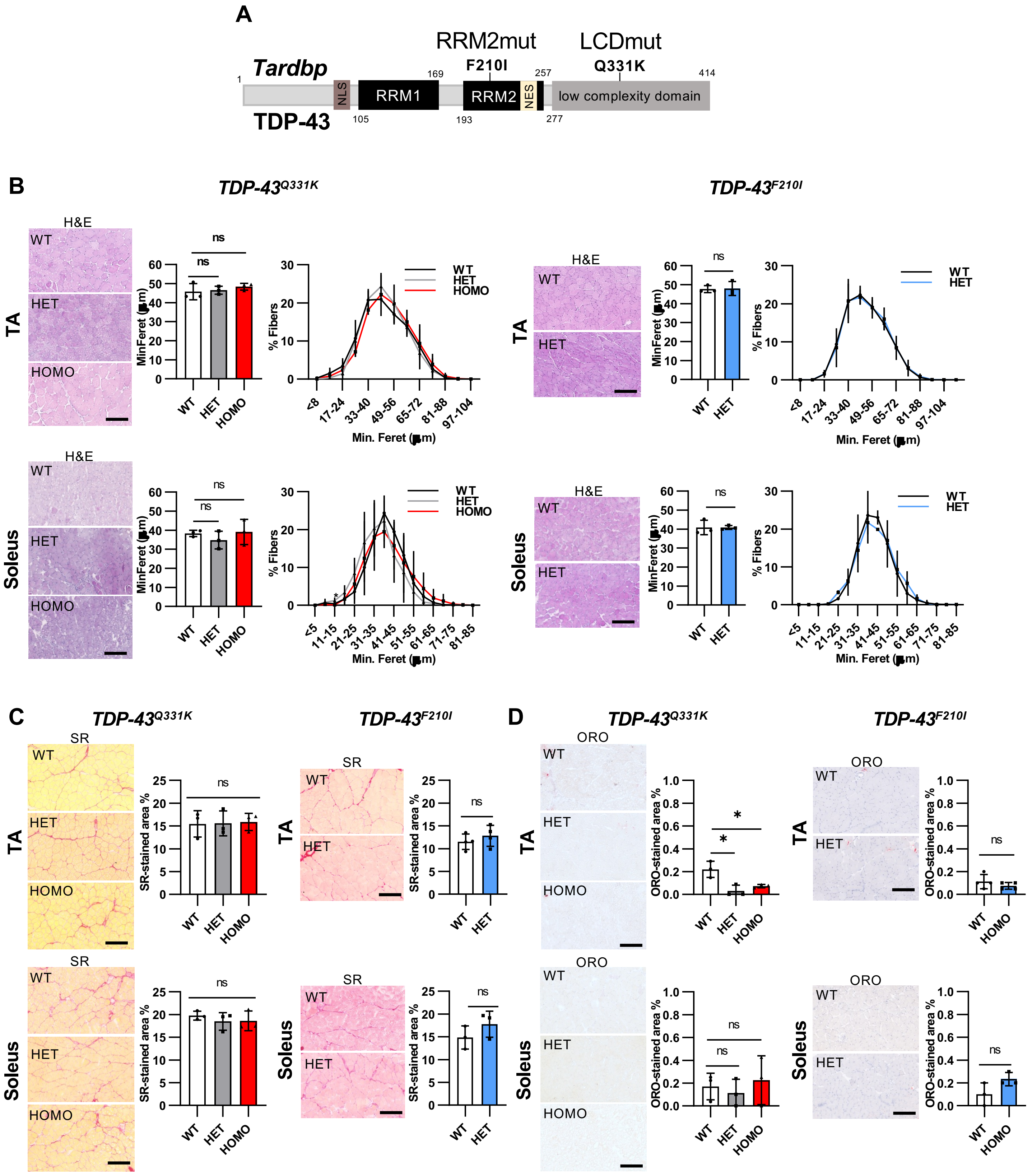

### Supplementary Figure 2

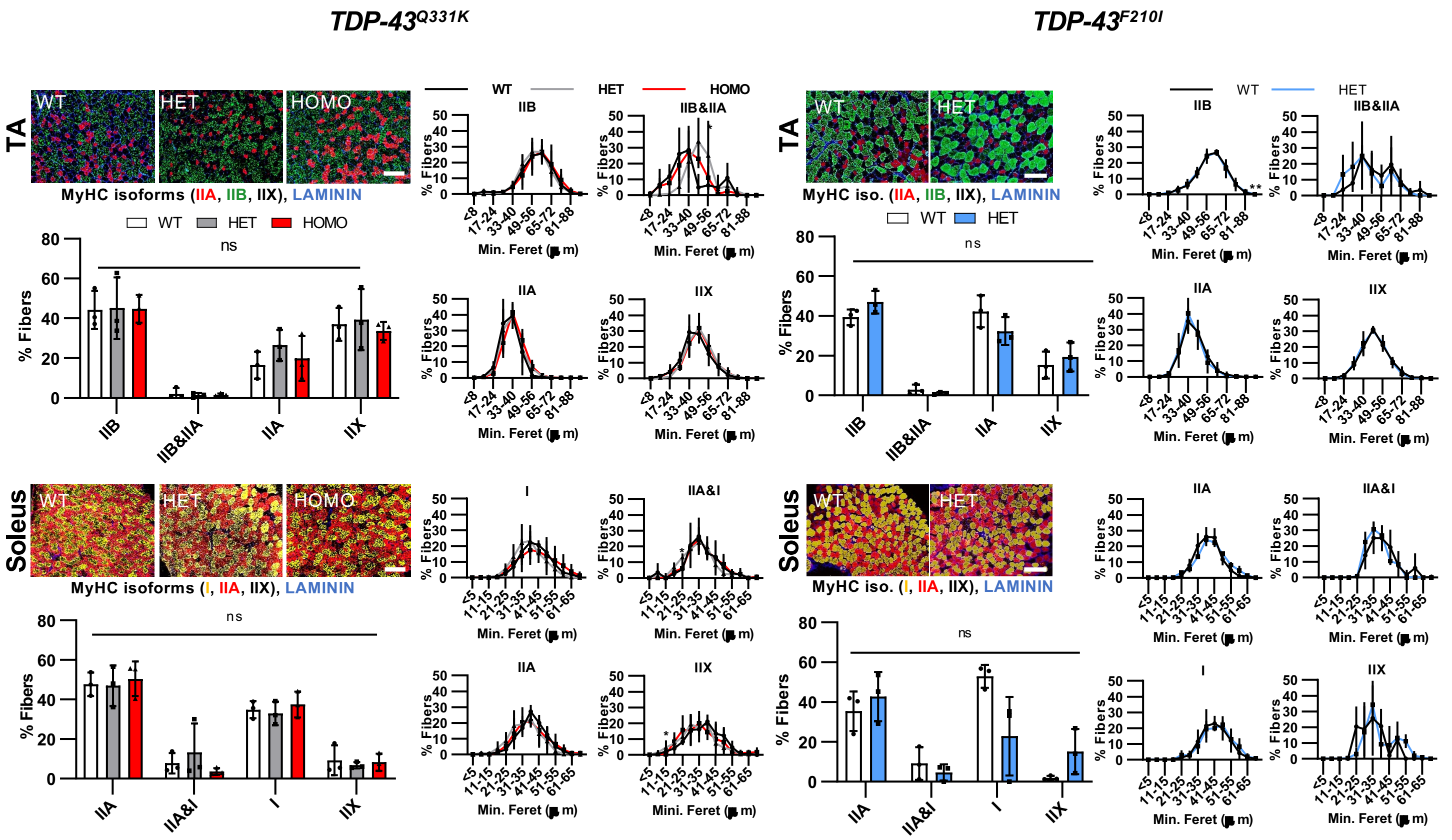

### Supplementary Figure 3

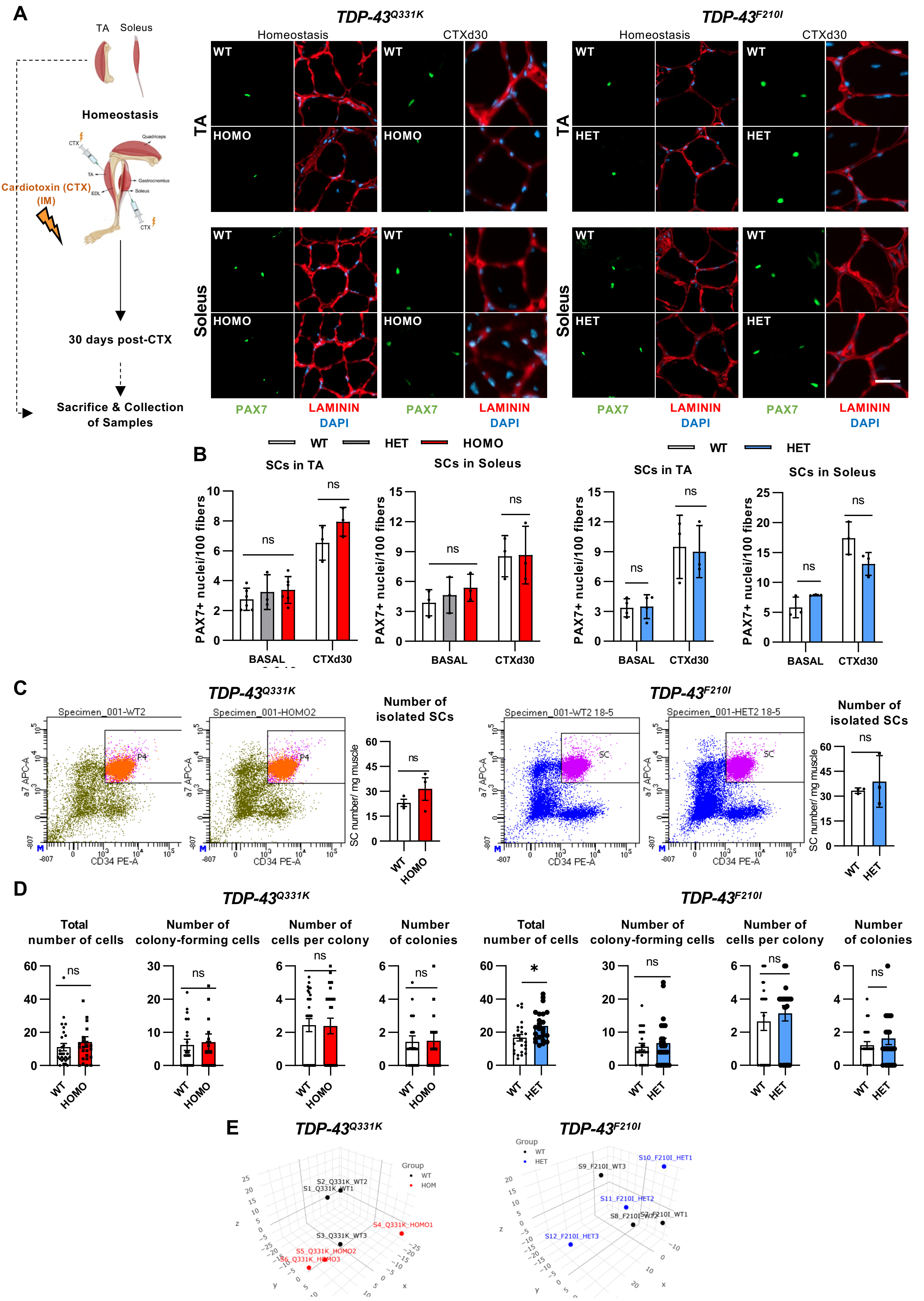

### Supplementary Figure 4

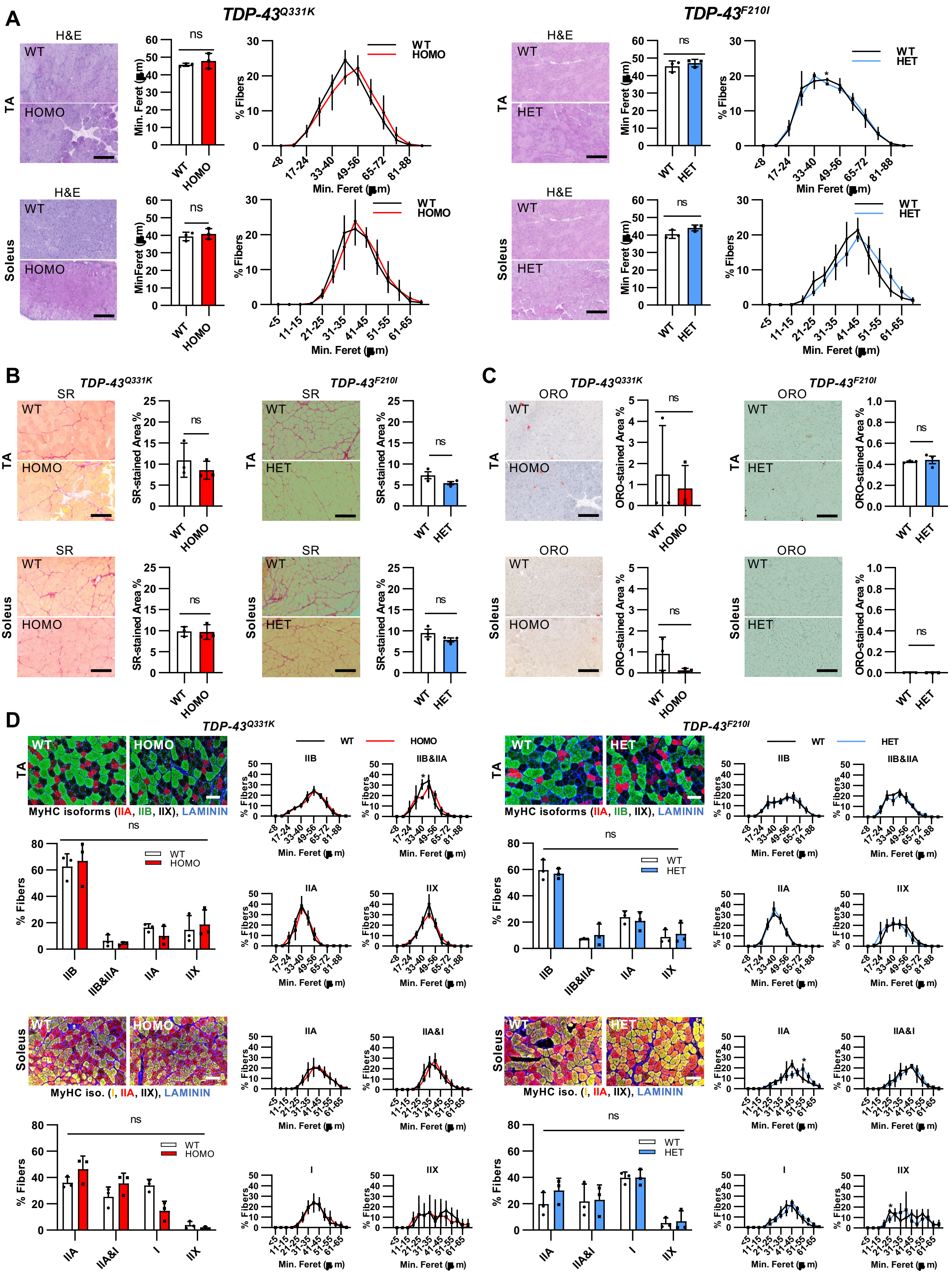

### Supplementary Figure 5

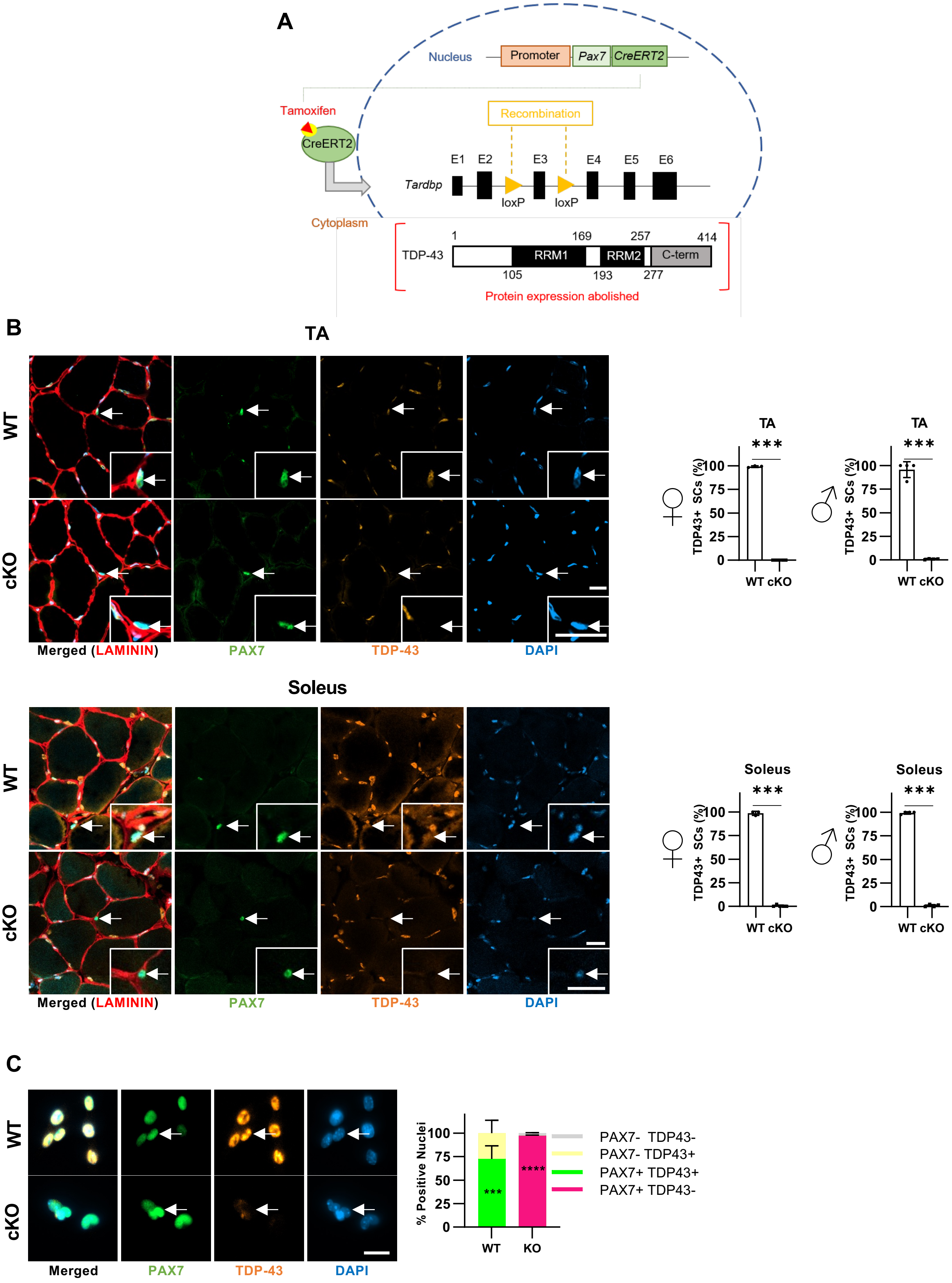

### Supplementary Figure 6

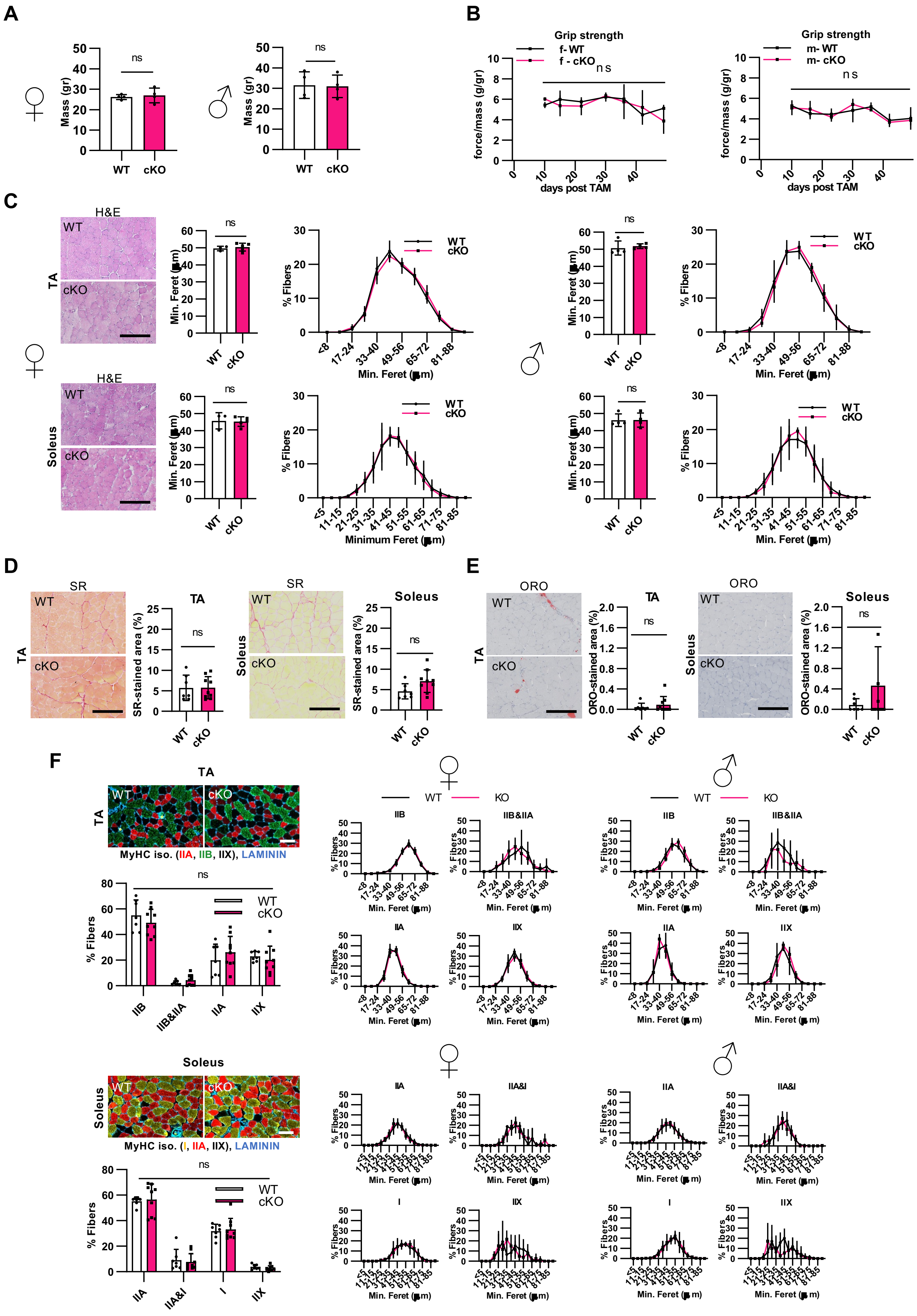

### Supplementary Figure 7

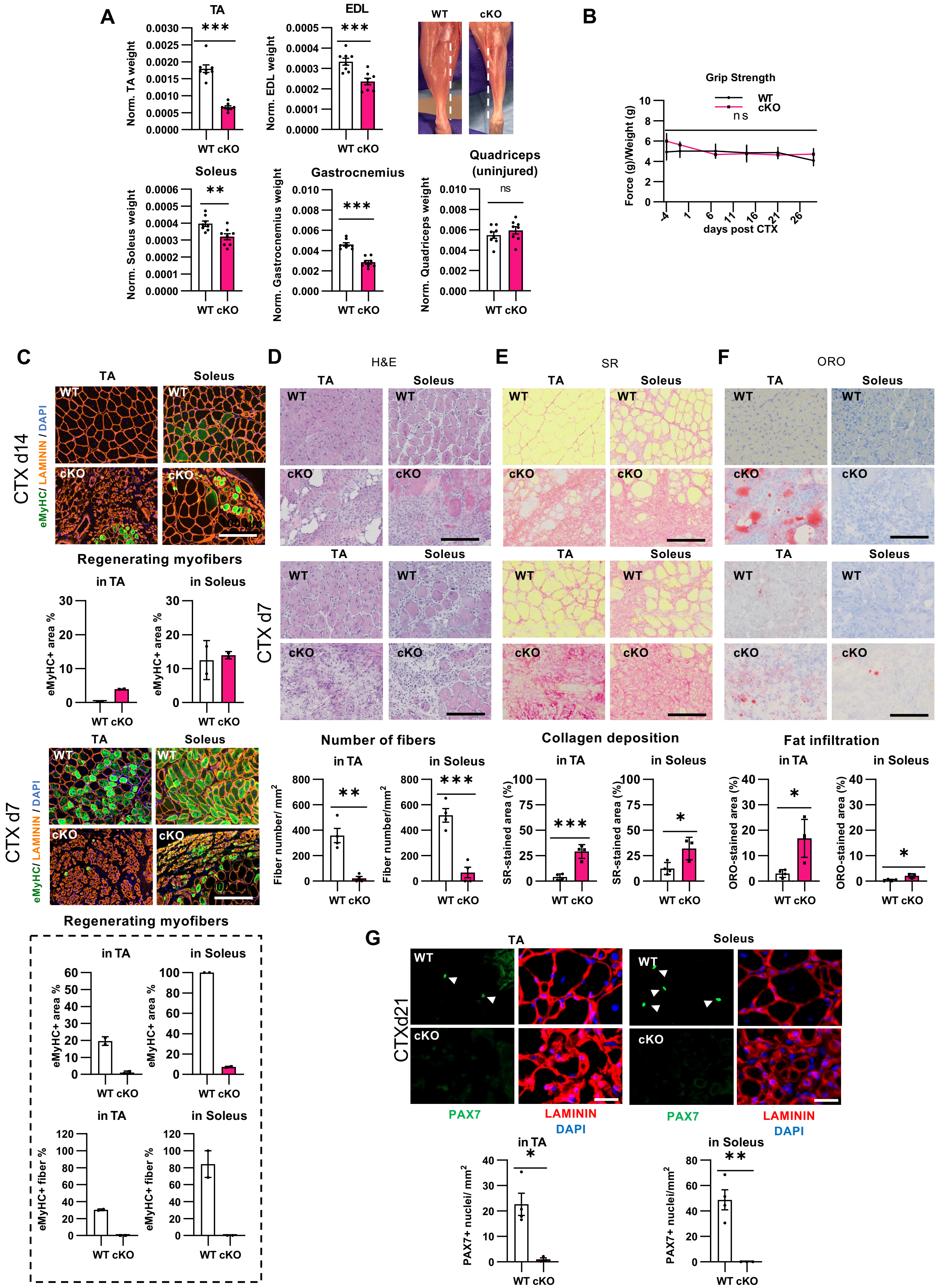

### Supplementary Figure 8

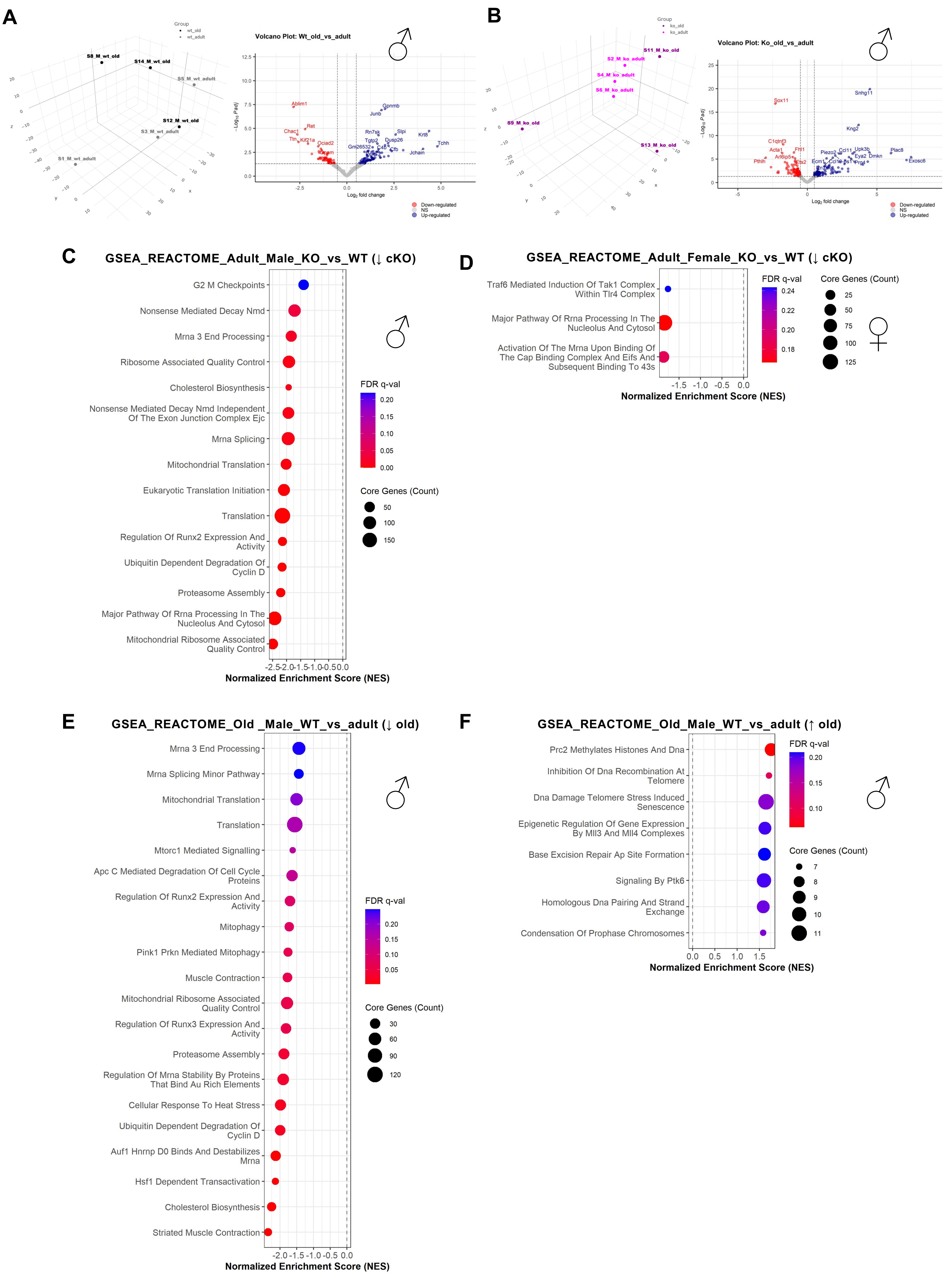

### Supplementary Figure 9

A

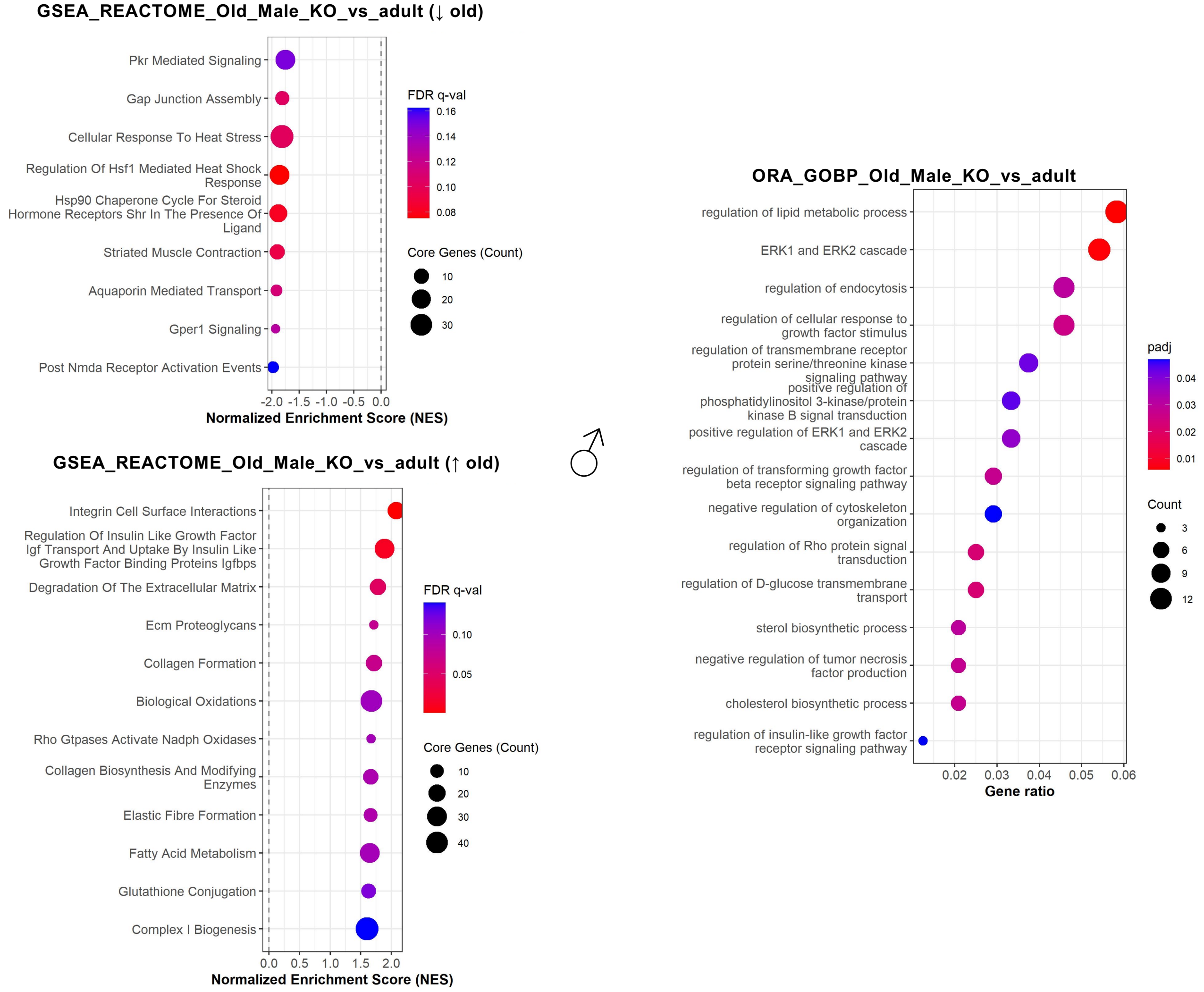


B

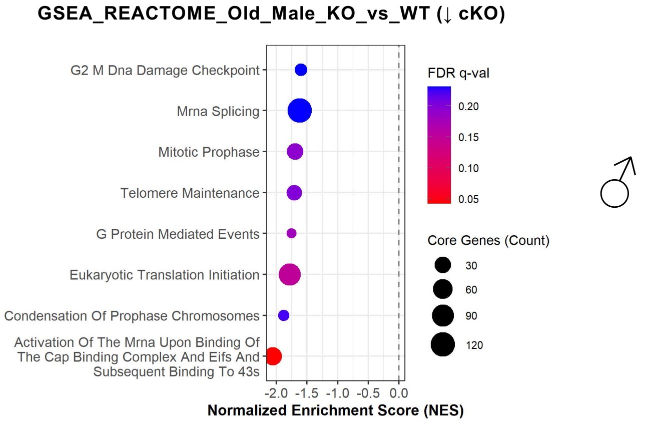

### Supplementary Figure 10

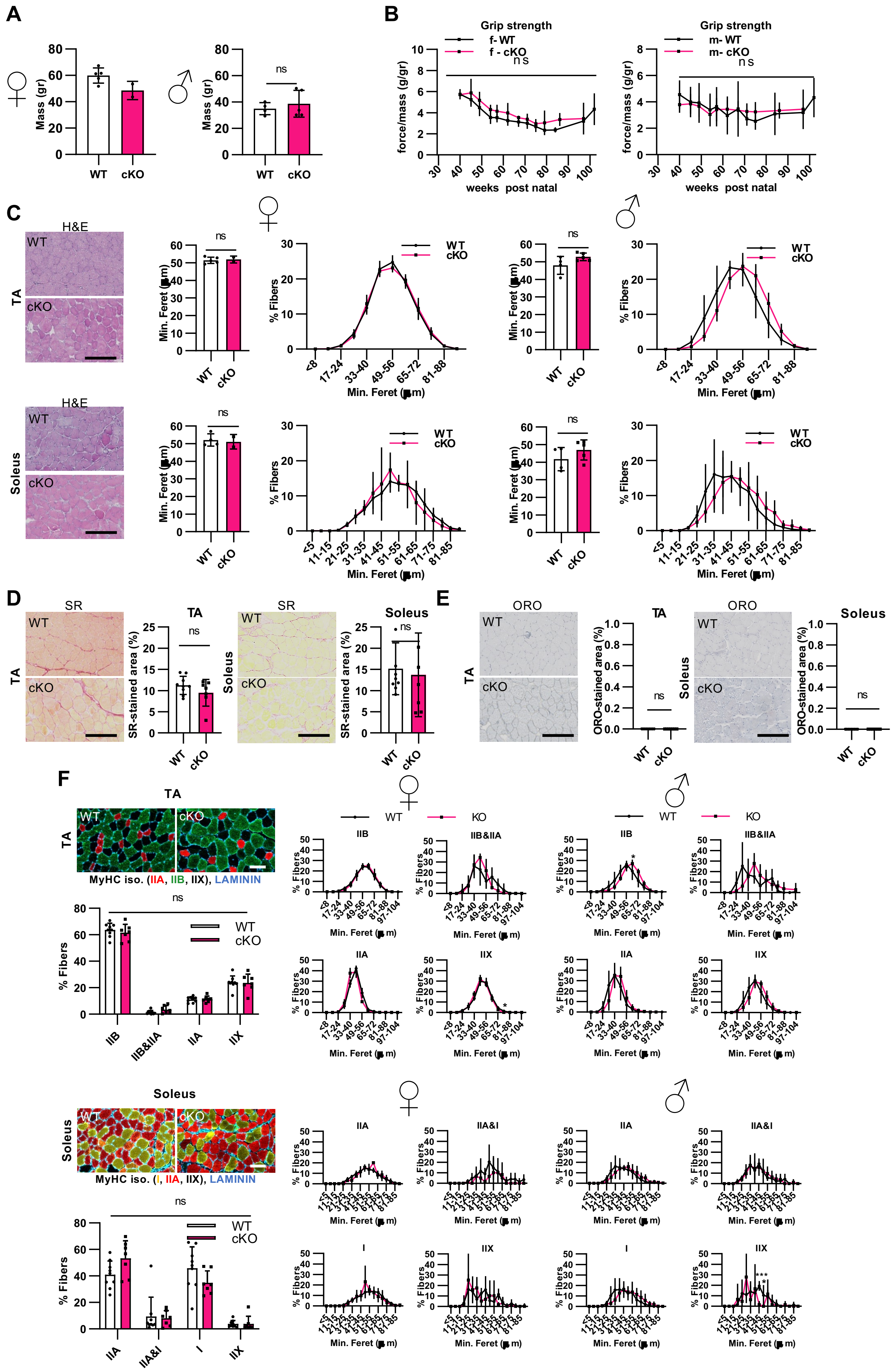

**Supplementary Figure 1. TDP-43 gain-of-function Q331K and loss-of-function F210I point mutations do not alter skeletal muscle homeostasis. (A)** Location of the Q331K and F210I point mutations within the *Tardbp* gene in the analysed TDP-43 knock-in mouse models. The F210I mutation falls within the RNA Recognition Motif 2 (RRM2), while Q331K mutation is located in the prion-like Low Complexity Domain (LCD) of TDP-43. Numbers indicate amino-acid residues of the TDP-43 protein encoded by *Tardbp*. NES: nuclear export signal. NLS: nuclear localization signal. **(B)** Representative images of Haematoxylin and Eosin (H&E) staining, along with histograms of mean myofiber size (CSA) and size distribution, in cross-sections of *tibialis anterior* (TA, top) and soleus (bottom) muscles from adult heterozygous (HET) and homozygous (HOMO) *TDP-43*$^{Q331K}$ mice (left panel) and adult HET *TDP-43*$^{F210I}$ mice (right panel), compared with their control (WT) littermates. **(C-D)** Representative images and quantification of the percentage of muscle area stained by Sirius Red (SR) for collagen deposition **(C)**, and Oil Red O (ORO) for fat infiltration **(D)** in TA (top) and soleus (bottom) muscles from HET and HOMO *TDP-43*$^{Q331K}$ mice (left panel) and HET *TDP-43*$^{F210I}$ mice (right panel), compared with their WT littermates. Bar graphs represent mean ± SD. N=3 per condition. Student's t test. *$p$<0.05; ns, not significant. Scale bars: 200 μm.

**Supplementary Figure 2. Skeletal muscle myofiber composition is preserved in *TDP-43*$^{Q331K}$ and *TDP-43*$^{F210I}$ mice**. Representative images and corresponding histograms showing the percentage of fiber isoforms and their CSA, determined by immunolabelling of myosin heavy chain (MyHC) isoforms: type IIA (red), IIB (TA-sections, green), IIX (unlabelled-black) and type I (soleus sections, yellow), and LAMININ (basal lamina, blue) in TA (top) and soleus (bottom) muscles from HET and HOMO *TDP-43*$^{Q331K}$ mice (left panel) and HET *TDP-43*$^{F210I}$ mice (right panel), compared with their WT littermates. Bar graphs represent mean ± SD. N=3 per condition. Two-way ANOVA (left) and Student's t test per bin (right). ns, not significant. Scale bars: 200 μm.

**Supplementary Figure 3. Satellite cell homeostasis and regeneration are preserved in *TDP-43*$^{Q331K}$ and *TDP-43*$^{F210I}$ mice**. **(A)** Left: Experimental scheme for cardiotoxin (CTX) injury. Right: Representative cross-sectional images of *tibialis anterior* (TA, top) and soleus (bottom) muscles from adult heterozygous (HET) and homozygous (HOMO) *TDP-43*$^{Q331K}$ mice (left panel) and adult HET *TDP-43*$^{F210I}$ mice (right panel), compared with their control (WT) littermates, immunolabelled for PAX7 (satellite cells-SC, green), LAMININ (basal lamina, red) and nuclei with DNA intercalant DAPI (blue). **(B)** Quantification of PAX7+ nuclei (SCs) per 100 fibers. **(C)** Representative plots od FACS gating and quantification of SC numbers per mg tissue in uninjured muscles. **(D)** Clonogenic capacity of FACS-isolated SCs; dots represent individual wells. *TDP-43*$^{Q331K}$ adult males and *TDP-43*$^{F210I}$ adult females. Bar graphs represent mean ± SD. N=3. One-way ANOVA was performed to compare all groups (**B**, N=3), and pairwise comparisons were done using Student's t test. *$p$<0.05; ns, not significant. **(E)** Principal Component Analysis (PCA) plots for the RNA-seq assay performed with isolated SCs from *TDP-43*$^{Q331K}$ and *TDP-43*$^{F210I}$ mice. Scale bars: 50 μm.

**Supplementary Figure 4. Skeletal muscle regeneration is preserved in *TDP-43*$^{Q331K}$ and *TDP-43*$^{F210I}$ mice following injury. (A)** Representative images of Haematoxylin and Eosin (H&E) staining, along with histograms of mean myofiber size (CSA) and size distribution, in cross-sections of *tibialis anterior* (TA, top) and soleus (bottom) muscles from adult homozygous (HOMO) *TDP-43*$^{Q331K}$ mice (left panel) and adult heterozygous (HET) *TDP-43*$^{F210I}$ mice (right panel), compared with their control (WT) littermates, 30 days (d) after cardiotoxin (CTX) injury. **(B-C)** Representative images and quantification of the percentage of muscle area stained by Sirius Red (SR) for collagen deposition **(B)**, and Oil Red O (ORO) for fat infiltration **(C)** in TA (top) and soleus (bottom) muscles from HOMO *TDP-43*$^{Q331K}$ mice (left panel) and HET *TDP-43*$^{F210I}$ mice (right panel), compared with their WT littermates at CTX d30. **(D)** Representative images and corresponding histograms showing the percentage of fiber isoforms and their CSA, determined by immunolabelling of myosin heavy chain (MyHC) isoforms: type IIA (red), IIB (TA-sections, green), IIX (unlabelled-black) and type I (soleus sections, yellow), and LAMININ (basal lamina, blue) in TA (top) and soleus (bottom) muscles from HOMO *TDP-43*$^{Q331K}$ mice (left panel) and HET *TDP-43*$^{F210I}$ mice (right panel), compared with their WT littermates at CTX d30. *TDP-43*$^{Q331K}$ adult males and *TDP-43*$^{F210I}$ adult females. Bar graphs represent mean ± SD. N=3. Two-way ANOVA (left) and Student's t test per bin (right). ns, not significant. Scale bars: 200 μm.

**Supplementary Figure 5. TDP-43 depletion in satellite cells from adult *Pax7*$^{CreERT2/+}$*;Tardbp*$^{fl/fl}$ mice.** **(A)** Schematic representation of the conditional gene targeting strategy. Upon tamoxifen (TAM) administration, CreERT2 translocates to the nucleus and mediates recombination between *loxP* sites, flanking the exon 3 of the *Tardbp* gene encoding TDP-43 protein, to delete it. **(B)** Representative images of *tibialis anterior* (TA) and soleus cross-sections immunolabelled for PAX7 (satellite cells-SC, green), TDP-43 (yellow), LAMININ (basal lamina, red), and nuclei counter-stained with DAPI (blue) from *Pax7*$^{CreERT2/+}$*;Tardbp*$^{fl/fl}$ (cKO) in comparison to *Pax7*$^{+/+}$*;Tardbp*$^{fl/fl}$ control (WT) mice 60 days (d60) post tamoxifen (TAM) injection. Scale bars: 25 μm. **(D)** Isolated SCs after 72 hours (h) in culture, immunolabelled for PAX7 (green), TDP-43 (yellow), and DAPI (blue), from cKO and WT mice. Scale bar: 20 μm. White arrows indicate PAX7+ SCs. Bar graphs represent mean ± SD. N=4. Student's t test. ****p*<0.001; *****p*<0.0001.

**Supplementary Figure 6. TDP-43 depletion in satellite cells does not alter adult skeletal muscle homeostasis.** **(A)** Body weight of adult *Pax7*$^{CreERT2/+}$;*Tardbp*$^{fl/fl}$ (cKO) mice and *Pax7*$^{+/+}$; *Tardbp*$^{fl/fl}$ control (WT) littermates, females and males, respectively. **(B)** Grip strength normalized to body weight. f-, female; m-, male. **(C)** Representative images of Haematoxylin and Eosin (H&E) staining, along with histograms of mean myofiber size (CSA) and size distribution, in cross-sections of *tibialis anterior* (TA, top) and soleus (bottom) muscles from adult cKO female (left) and male (right) mice, compared with WT littermates. **(D-E)** Representative images (female TA and soleus) and quantification of the percentage of muscle area stained by Sirius Red (SR) for collagen deposition **(D)**, and Oil Red O (ORO) for fat infiltration **(E)** in TA and soleus muscles from adult cKO female (left) and male (right) mice, compared with WT littermates. **(F)** Representative images and corresponding histograms showing the percentage of fiber isoforms and their CSA, determined by immunolabelling of myosin heavy chain (MyHC) isoforms: type IIA (red), IIB (TA-sections, green), IIX (unlabelled-black) and type I (soleus-sections, yellow), and LAMININ (basal lamina, blue) in TA (female, top) and soleus (female, bottom) muscles from adult cKO female (left) and male (right) mice, compared with WT littermates. Bar graphs represent mean ± SD. Student's t test **(A-E)**, two-way ANOVA (left) and Student's t test per bin (right) **(F)**. ns, not significant. Scale bars: 200 μm.

**Supplementary Figure 7. TDP-43 depletion in female satellite cells abolishes muscle regeneration and stem cell pool restoration upon cardiotoxin injury. (A)** As in **Figure 5A-B** for males, *Tibialis anterior* (TA), soleus, *extensor digitorum longus* (EDL), gastrocnemius and quadriceps muscle weights from female *Pax7*$^{CreERT2/+}$;*Tardbp*$^{fl/fl}$ (cKO) mice and *Pax7*$^{+/+}$;*Tardbp*$^{fl/fl}$ control (WT) littermates, 30 days (d) post cardiotoxin-injury (CTX d30), and hindlimb photographs illustrating muscle loss. Dotted lines indicate tibia length (~1.7 cm). Bar graphs represent mean ± SD. N=8. **(B)** Grip strength normalized to body weight over the d7-14-21-30 time-points post-CTX. **(C-F)** Top: representative images; bottom: quantification of TA and soleus muscles at CTX d14 and CTX d7 stained with **(C)** embryonic myosin heavy chain (eMyHC, newly formed myofibers, green), LAMININ (basal lamina, orange), and DAPI (nuclei, blue) and quantification of eMyHC+ area; **(D)** Haematoxylin & Eosin (H&E) for morphology and quantification of number of fibers per area; **(E)** Sirius Red (SR) for collagen deposition (area quantified); and **(F)** Oil Red O (ORO) for fat infiltration (area quantified). Bar graphs represent mean ± SD. N=2 per condition. **(G)** Upper panel: representative cross-sectional images of TA and soleus muscles from female WT and cKO mice at CTX d21 immunolabelled for PAX7 (satellite cells-SC, green), LAMININ (red) and DAPI (blue). Lower panel: quantification of PAX7+ SCs in TA and soleus muscles at CTXd 14-21. Arrowheads indicate SCs. Scale bars: Bar graphs represent mean ± SD. N=1-2 per condition. Student's t test. *$p$<0.05; **$p$<0.01; ***$p$<0.001.

**Supplementary Figure 8. TDP-43 depletion remodels the transcriptomic profiles of adult satellite cells resembling signatures of physiological aging. (A-B)** Principal Component Analysis (PCA) of RNA-seq and Volcano Plot of differentially expressed genes (DEG) ($|\log_2 FC| \geq 0.5$, adjusted $p$<0.05) for satellite cells (SC) from adult and old *Pax7*$^{+/+}$;*Tardbp*$^{fl/fl}$ control (WT) **(A)** and *Pax7*$^{CreERT2/+}$;*Tardbp*$^{fl/fl}$ (cKO) **(B)** male mice. **(C-D)** Gene Set Enrichment Analysis (GSEA) results against the Reactome dataset (FDR q < 0.25) for adult from cKO and control WT male **(C)** and female **(D)** mice. **(E-F)** GSEA results against the Reactome dataset (FDR q < 0.25) for old and adult WT male mice, showing downregulated **(E)** and upregulated **(F)** pathways for SCs from old mice. N=3.

**Supplementary Figure 9. Aging produces limited additional transcriptional changes in TDP-43-deficient satellite cells. (A)** Gene Set Enrichment Analysis (GSEA) results against the Reactome dataset (FDR q < 0.25) for satellite cells (SC) from adult and old *Pax7*$^{CreERT2/+}$;*Tardbp*$^{fl/fl}$ (cKO) male mice. **(B)** GSEA results against the Reactome dataset (FDR q < 0.25) for satellite cells from old cKO and *Pax7*$^{+/+}$;*Tardbp*$^{fl/fl}$ control WT male mice. N=3.

**Supplementary Figure 10. TDP-43 depletion in satellite cells does not alter skeletal muscle integrity in aged mice. (A)** Body weight of aged *Pax7*$^{CreERT2/+}$;*Tardbp*$^{fl/fl}$ (cKO) mice and *Pax7*$^{+/+}$; *Tardbp*$^{fl/fl}$ control (WT) littermates, females and males, respectively. **(B)** Grip strength normalized to body weight over the life-time of aged cKO and WT mice post-tamoxifen (TAM) administration. **(C)** Representative images of Haematoxylin and Eosin (H&E) staining, along with histograms of mean myofiber size (CSA) and size distribution, in cross-sections of *tibialis anterior* (TA, top) and soleus (bottom) muscles from aged cKO female (left) and male (right) mice, compared with WT littermates. **(D-E)** Representative images (female TA and soleus) and quantification of the percentage of muscle area stained by Sirius Red (SR) for collagen deposition **(D)**, and Oil Red O (ORO) for fat infiltration **(E)** in TA and soleus muscles from aged cKO female (left) and male (right) mice, compared with WT littermates. **(F)** Representative images and corresponding histograms showing the percentage of fiber isoforms and their CSA, determined by immunolabelling of myosin heavy chain (MyHC) isoforms: type IIA (red), IIB (TA-sections, green), IIX (unlabelled-black) and type I (soleus-sections, yellow), and LAMININ (basal lamina, blue) in TA (female, top) and soleus (female, bottom) muscles from aged cKO female (left) and male (right) mice, compared with WT littermates. Bar graphs represent mean ± SD. Student's t test **(A-E)**, two-way ANOVA (left) and Student's t test per bin (right) **(F)**. $p<0.05$; ns, not significant.
